# A haplotype-resolved, chromosome-scale genome assembly and annotation for *Carya glabra* (pignut hickory; Juglandaceae)

**DOI:** 10.64898/2025.12.19.695579

**Authors:** Shengchen Shan, Edgardo M. Ortiz, Baylee Klein, Arthur Oganisyan, Gia Serrano, Bryanna Stults, Rubina Torkzadeh, Audrey Tucker, Ezra Linnan, Benjamin Pringle, Tyler Radtke, Moumita Hoque Rainy, Lia Swanson, Gabrielle Vines, Lauren Whitt, Huiting Zhang, Alex Harkess, Pamela S. Soltis, Douglas E. Soltis

## Abstract

*Carya glabra* (2*n* = 4*x* = 64), also known as pignut hickory, is a widely distributed species in the walnut family (Juglandaceae). Native to the central and eastern United States and southeastern Canada, *C. glabra* plays an important ecological role as a common upland forest species; it is closely related to several economically valuable nut trees, including *C. illinoinensis* (pecan). A deeper understanding of the genetics of *C. glabra* is essential for studying its evolutionary history and biology, with potential implications for agricultural improvement of pecan. Here, we present the first nuclear genome assembly and annotation of *C. glabra*. The assembly is chromosome-level and phased, representing the first assembled polyploid genome in the genus *Carya*. A total of 64 pseudochromosomes were assembled and phased into four haplotypes. The haplotype A assembly spans 600.4 Mb, comprises 55.0% repetitive sequences, and contains 30,947 protein-coding genes, with a BUSCO completeness score of 97.7%. Functional annotation assigned 94.3% of haplotype A genes to gene families, and 79.7% and 86.3% of genes were annotated with Gene Ontology terms and protein domains, respectively; 635 putative plant disease resistance genes were found in haplotype A. The other three haplotypes exhibited similarly high-quality annotation metrics. Our genomic analyses also suggest that *C. glabra* is an autotetraploid. Comparative genomic analyses revealed high collinearity among the four haplotypes of *C. glabra* and the published genomes of three other *Carya* species, although structural variation among the genomes of these species was identified. In addition, we provide an improved chloroplast genome assembly and the first mitochondrial genome for *C. glabra*. Importantly, most members of the research team are undergraduate students; the sequenced individual is located in McCarty Woods, a Conservation Area on the University of Florida campus. This work highlights the value of genome assembly efforts as powerful tools for teaching genomics and supporting conservation initiatives. This first high-quality reference genome for *C. glabra* provides a valuable resource for studying *Carya*, a genus of significant ecological and economic importance.

**Article summary:** *Carya glabra* (pignut hickory) is a common upland forest species in North America. This species is a member of the walnut family (Juglandaceae), which includes many economically important nut trees. Here, we present the first nuclear genome assembly and annotation of *C. glabra*. The assembly is chromosome-level and phased. The haplotype A assembly contains 30,947 protein-coding genes, with a BUSCO completeness score of 97.7%. Our genomic analyses suggest that *C. glabra* is an autopolyploid. We also provide chloroplast and mitochondrial genome assemblies. This nuclear genome provides a valuable resource for studying *Carya*, a genus of significant ecological and economic importance.

## Introduction

*Carya glabra* (2*n* = 4*x* = 64) (Juglandaceae; walnut family), commonly known as pignut hickory, is a widespread species in the central and eastern United States and southeastern Canada, ranging from Ontario southward to central Florida (Fig. 1a; POWO 2025). Pignut hickory is a slow-growing, deciduous tree that typically reaches 20–30 meters in height and 30–100 centimeters in diameter (Tirmenstein 1991). The species is monoecious, bearing staminate catkins and pistillate flowers that appear in spikes (Tirmenstein 1991). *Carya* possesses an accessory fruit; a pear-shaped nut is enclosed in a four-valved husk (of bracts). The fruit remains green until maturity, turning brown as it ripens (Fig. 1a; Smalley 1990).

**Fig. 1.**
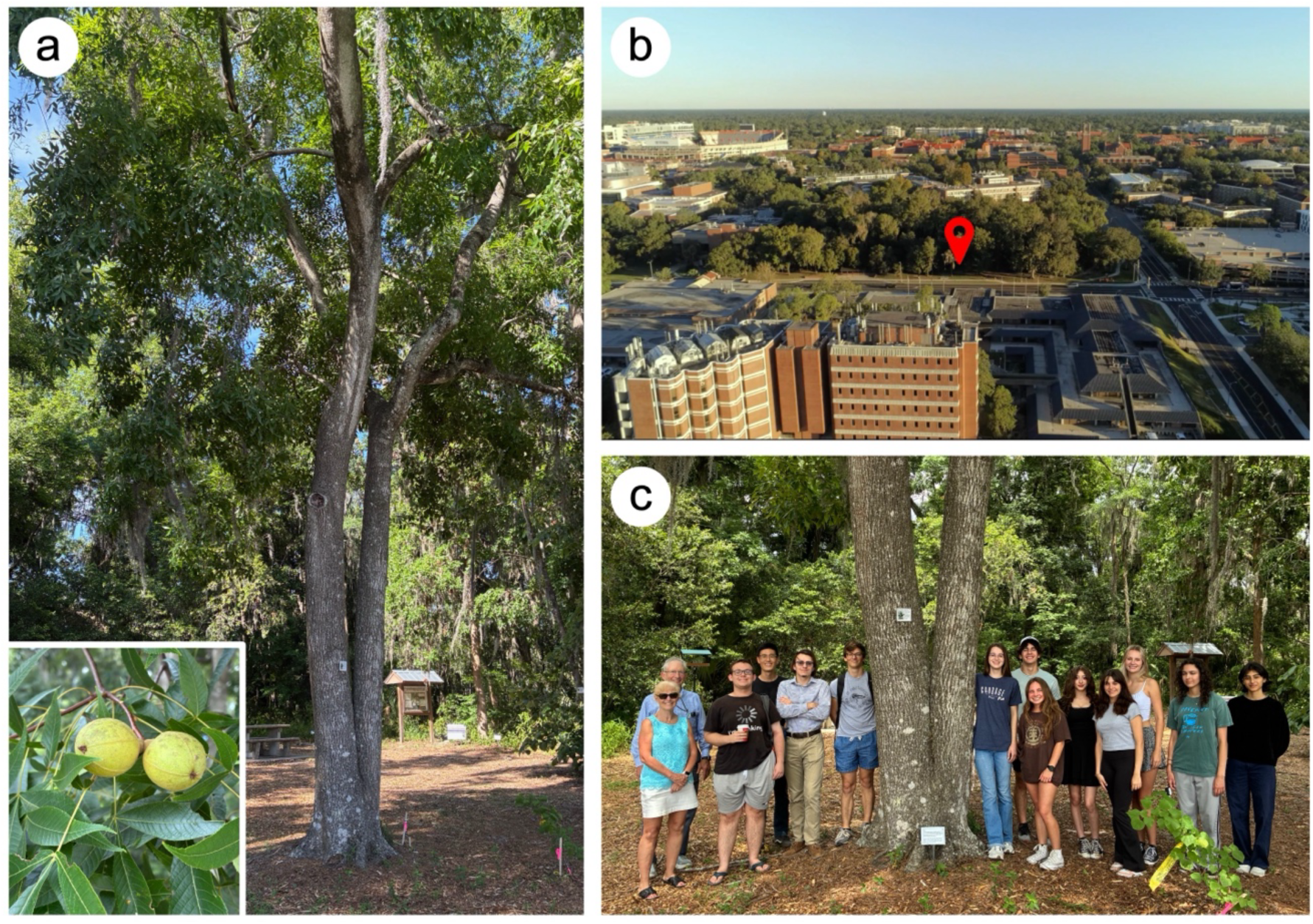
*Carya glabra* (pignut hickory) on the campus of the University of Florida. (a) The *C. glabra* individual sequenced in this study; the inset highlights the fruits and compound leaves. (b) Location of the *C. glabra* individual (indicated by the red pin) in McCarty Woods on the University of Florida campus. (c) Most members of the research team in front of the *C. glabra* tree; most are undergraduate researchers. Photo credits: (a) Shengchen Shan; (b) John Rouse; (c) Erin L. Grady.

The species is an ecological dominant in dry upland forests (Smalley 1990). In addition, the nuts are rich in crude fat and are consumed by a variety of wildlife, including squirrels, birds, foxes, rabbits, and raccoons (Smalley 1990). The wood of *C. glabra* is heavy and strong, making it ideal for tool handles and mallets, and it is also commonly used as fuelwood (Smalley 1990; Tirmenstein 1991). Pignut hickory also shows potential value for restoration of disturbed sites, as it has been reported to recolonize abandoned strip mines (Hardt and Forman 1989).

*Carya* comprises 19 species with an intercontinentally disjunct distribution (POWO 2025). In Asia, the genus is native to India, China, and countries in Southeast Asia, while in North America it occurs in eastern Canada, central and eastern United States, and Mexico (POWO 2025). Phylogenetic analyses support two monophyletic groups within the genus, corresponding to the primary geographic distributions (Asia and North America) (Zhang et al. 2013; Xi et al. 2022; Zhang et al. 2024b). According to molecular age estimation and biogeographic analyses, *Carya* in North America dates to the early Paleocene (Zhang et al. 2013). Its earliest confirmed occurrence is evidenced by fossil fruits from the late Eocene (Manchester 1999). The highest species diversification rate of the North America clade occurred around 10.1 million years ago (Ma) during the late Miocene, suggesting that *C. glabra* or its ancestor likely emerged around this time (Zhang et al. 2013). At least six North American *Carya* species, including *C. glabra*, are tetraploid (2*n* = 4*x* = 64) (Woodworth 1930; Stone 1961; Zhang et al 2013), whereas all Asian species investigated are diploid (Grauke 2016). The North America clade showed a higher diversification rate than the Asia clade, which may be attributed to the polyploid nature of many North American species (Zhang et al. 2013).

Recent phylogenetic studies indicate that the closest relative of *C. glabra* may be *C. texana*, which is also a tetraploid (Huang et al. 2019; Xi et al. 2022). Based on plastome data, other close relatives include *C. palmeri* (2*n* = 2*x* = 32) and some but not all populations of *C. illinoinensis* (2*n* = 2*x* = 32) (Xi et al. 2022). In contrast, phylogenetic analyses, using approximately 10× resequencing data relative to the *C. cathayensis* genome, indicate that the clade containing *C. glabra* and *C. texana* is sister to another tetraploid species, *C. tomentosa* (Huang et al. 2019). Notable reported examples of natural hybridization involving *C. glabra* include the hybrid *Carya* × *demareei* Palmer, which arose from a cross between *C. glabra* and diploid *C. cordiformis* (Sutton and Crowley 2020). Furthermore, the overlapping geographical ranges of *C. glabra* and tetraploid *C. ovalis* have led to frequent hybridization between those two species (Coder 2023).

*Carya* includes two species that are commercially cultivated nut trees: *C. illinoinensis* (pecan) and *C. cathayensis* (Chinese hickory) (Grauke 2016). In the United States, pecan production exceeded 120,000 metric tons in 2024, with a value of $468 million (USDA-NASS 2025). To date, genome assemblies have been reported for three *Carya* species – *C. illinoinensis* (Huang et al. 2019; Lovell et al. 2021; Xiao et al. 2021), *C. cathayensis* (Huang et al. 2019; Zhang et al. 2024b), and *C. sinensis* (Zhang et al. 2024b) – all of which are diploid.

In this study, we assembled and annotated the first nuclear genome of tetraploid *Carya glabra*. This chromosome-level, phased genome represents the first polyploid genome reported within the genus. The reference genome of *C. glabra* should enable novel research in the economically important genus *Carya*, with broad applications in both agriculture and evolutionary biology. The sequenced individual is located in McCarty Woods, a designated Conservation Area and quiet oasis at the center of the University of Florida (UF) campus (Fig. 1b). Most of the researchers involved in this project are undergraduate students enrolled in a Course-based Undergraduate Research Experience (CURE) class at UF (Fig. 1c). As part of the American Campus Tree Genomes (ACTG) project (https://www.hudsonalpha.org/actg), this work highlights the potential of genome assembly projects to support conservation efforts and enhance hands-on genomics education.

## Materials & Methods

### Sample collection

Fresh leaf and axillary bud tissues were collected from a *Carya glabra* individual in the McCarty Woods Conservation Area, located centrally on the UF campus. An herbarium voucher for this plant was deposited in the Florida Museum of Natural History Herbarium (FLAS). The collected tissues were immediately frozen in liquid nitrogen.

### DNA isolation and sequencing

*Carya glabra* leaf tissue was sent to the HudsonAlpha Institute for Biotechnology (Huntsville, AL, USA) for DNA isolation and subsequent sequencing. High-molecular-weight DNA was extracted using the Nanobind Plant Nuclei Big DNA Kit (Circulomics-PacBio, Menlo Park, CA, USA). Isolated DNA was sheared with Megaruptor (Diagenode, Denville, NJ, USA), and fragments with a size of approximately 25 kb were selected using BluePippin (Sage Science, Beverly, MA, USA). Size-selected DNA was used to construct the PacBio sequencing library using the SMRTbell Express Template Prep Kit 2.0 (PacBio, Menlo Park, CA, USA). The library was then sequenced on two SMRT Cells on a PacBio Revio system at HudsonAlpha to generate High-Fidelity (HiFi) reads.

In addition, an Omni-C library was constructed using flash-frozen leaf material following the Dovetail Genomics protocol (Dovetail Genomics, Scotts Valley, CA, USA). The library was sequenced on one S4 flow cell of the Illumina NovaSeq 6000 system (Illumina, San Diego, CA, USA) at HudsonAlpha to generate paired-end 150-bp reads. Basic statistics of PacBio HiFi data and Omni-C data were assessed using SeqKit2 (v.2.4.0; Shen et al. 2024).

### RNA isolation and sequencing

Leaf and axillary bud tissues from the same *C. glabra* individual used for DNA isolation were collected and flash-frozen in liquid nitrogen. RNA was extracted from each tissue (leaf and axillary bud) using a modified CTAB method (Jordon-Thaden et al. 2015). RNA quality was assessed using a Bioanalyzer at the Interdisciplinary Center for Biotechnology Research (ICBR), UF (Gainesville, FL, USA). Two strand-specific (i.e., directional) RNA-seq libraries were prepared, and the libraries were sequenced on the Illumina NovaSeq X platform to generate paired-end 151-bp reads at ICBR. The statistics of the RNA-seq data were calculated using SeqKit2, and the raw reads were filtered using fastp (v.0.23.4; Chen et al. 2018) with default parameters.

### Chloroplast and mitochondrial genome assembly and annotation

Both organellar genomes were simultaneously assembled from PacBio HiFi reads using Oatk (v1.0; Zhou et al. 2025). Oatk’s plastome assembly graph was simplified and circularized using Bandage (v.0.8.1; Wick et al. 2015), and the resulting assembly was annotated using the web application GeSeq (https://chlorobox.mpimp-golm.mpg.de/geseq.html; Tillich et al. 2017).

The plastome annotation was further curated by comparing GeSeq’s annotation with the well-annotated *Nicotiana tabacum* chloroplast genome (NCBI accession number: NC_001879), as well as three published *Carya glabra* chloroplast genomes (BK061156; OR099205; NC_067504) (Luo et al. 2021; Xi et al. 2022; Liu et al. 2025). The chloroplast genomes were first aligned using MAFFT (v.7.490) with default parameters in Geneious Prime (2025.2.2; https://www.geneious.com). The annotation was then manually inspected and curated. Ambiguous transfer RNA (tRNA) annotations were further validated using BLAST searches in the PlantRNA 2.0 database (http://plantrna.ibmp.cnrs.fr/; Cognat et al. 2022).

Oatk’s mitochondrial assembly graph could not be resolved into a single circular chromosome without excluding graph segments. Therefore, two circular contigs were inferred from the graph and saved as separate chromosomes using Bandage. These two mitochondrial chromosomes were annotated with the web application PMGA (http://47.96.249.172:16084/annotate.html; Li et al. 2025) using the three databases available in the program. Additionally, we searched plastome and mitochondrial proteins using Captus (v.1.6.1; Ortiz et al. 2023). The four annotation tracks, one from Captus and three from PMGA (each corresponding to one of the three databases from PMGA), were checked against each other for consistency, retaining only the best annotation (i.e., that includes start and stop codons whenever possible, longest and/or most frequently observed) in case of discrepancies.

Following manual curation, the edited GenBank files were exported from Geneious Prime and then uploaded to OGDRAW (v.1.3.1; Greiner et al. 2019) to generate the final chloroplast and mitochondrial genome annotation maps using the default parameters (except checking the “tidy up annotation” box).

### Nuclear genome profiling

Jellyfish (v2.3.0; Marçais and Kingsford 2011) was used to count *k*-mers and generate a *k*-mer histogram (*k*-mer size: 21) from the HiFi reads. The *k*-mer histogram was then imported to GenomeScope 2.0 (http://genomescope.org/genomescope2.0/; Ranallo-Benavidez et al. 2020) to infer nuclear genome characteristics, including monoploid genome size and heterozygosity, with default parameters except setting ploidal level as 4.

### Nuclear genome assembly

Hifiasm (v.0.19.9; Cheng et al. 2021) was used to perform *de novo* assembly with default parameters. Both HiFi reads and Omni-C reads were used as input data. Given the polyploid nature of the *Carya glabra* genome, the unitig assembly from hifiasm, which contained the genomic information from all four haplotypes, was used for downstream analyses.

To scaffold the unitigs, first, bwa-mem2 (v.2.2.1; Vasimuddin et al. 2019) was used to align the Omni-C reads to the unitig assembly. The resulting alignments were then analyzed with the hic_qc pipeline from Phase Genomics (Seattle, WA, USA) to assess the overall quality of the Omni-C library. Then, YaHS (v.1.1; Zhou et al. 2023) was used to perform the scaffolding process with default parameters.

Next, using the Hi-C alignment file as input, the ‘juicer pre’ tool from YaHS and Juicer (v.1.22.01; Durand et al. 2016) were used to generate the Hi-C contact map. We then manually curated the assembly by examining the Hi-C contact map using Juicebox Assembly Tools (v.1.11.08; Dudchenko et al. 2018). Misjoin and inversion errors were manually corrected, and the orientation of chromosomes was also curated to match the published *Carya illinoinensis* genome (Lovell et al. 2021). After all edits, the final genome assembly was generated using the ‘juicer post’ tool from YaHS.

A dot plot was generated using the web application D-GENIES (https://dgenies.toulouse.inra.fr/; Cabanettes and Klopp 2018) to compare the *Carya illinoinensis* genome with the assembled *C. glabra* genome. To assign scaffolds to chromosomes, the *C. glabra* scaffolds were renamed according to their alignment with the *C. illinoinensis* chromosomes. The four copies of each chromosome in *C. glabra* were labeled A, B, C, and D in descending order of length. Each set of 16 chromosomes with the same label (e.g., Chr01A, Chr02A, …, Chr16A) was grouped and referred to as a haplotype (e.g., haplotype A). The 64 chromosomes were therefore assigned to four haplotypes (A, B, C, and D). It is important to note that this haplotype assignment is artificial and does not necessarily reflect a biological haplotype, since each haplotype set may represent a mixture of chromosomes originating from different gametes. For each haplotype set, genome completeness was estimated using benchmarking universal single-copy orthologs (BUSCO, v.5.3.0) with the eudicots_odb10 database (Manni et al. 2021).

### Nuclear genome annotation

To annotate repeat sequences, for each haplotype of the chromosome-level genome assembly, EDTA (v.2.1.0; Ou et al. 2019) was used for *de novo* transposable element (TE) annotation. Using the TE library generated by EDTA, RepeatMasker (v.4.1.7; Smit et al. 2013-2015) was used to identify additional repeat elements and to softmask the genome (with repeat elements written in lowercase).

For gene annotation, BRAKER3 (v.3.0.8; Gabriel et al. 2024) was used to predict protein-coding genes using the RNA-seq data from the leaf and axillary bud tissues from *C. glabra* and protein evidence from model species (Table S1). Various BRAKER3 parameter settings were tested using the haplotype A genome (Table S2). The setting that resulted in the highest BUSCO score (using the eudicots_odb10 database) was applied to annotate all other haplotypes (i.e., B, C, and D). After the initial annotation, gene models meeting any of the following criteria were filtered out using AGAT (v.1.4.2; Dainat 2022): (1) presence of a premature stop codon; (2) absence of a start and/or stop codon; or (3) an open reading frame (ORF) length of ≤100 amino acids or ≤50 amino acids. The genes were named in accordance with the guidelines proposed by Cannon et al. (2025).

Functional annotation was performed using the web application TRAPID 2.0 (Bucchini et al. 2021), with the PLAZA 4.5 dicots database (Van Bel et al. 2018) as the reference and the rosids clade selected for the similarity search. All parameters were set to default, except that “input sequences are CDS” was selected.

Lastly, Circos (v.0.69-9; Krzywinski et al. 2009) was used to visualize the genome and the associated genetic features, including gene and TE densities along the chromosomes.

### Comparative genomic analyses

Genome-level synteny analysis was performed using GENESPACE (v.1.3.1; Lovell et al. 2022) to compare the four *Carya glabra* haplotypes with chromosome-level genome assemblies from three other *Carya* species: *C. cathayensis* (Zhang et al. 2024b), *C. illinoinensis* (Lovell et al. 2021), and *C. sinensis* (Zhang et al. 2024b).

### Identification of putative disease resistance genes

Because disease resistance is a key trait for pecan improvement, plant disease resistance genes (*R* genes) in the *Carya glabra* genome were predicted using the DRAGO 2 pipeline (with default parameters) from the Plant Resistance Genes database (PRGdb 3.0) (Osuna-Cruz et al. 2018). Using the same pipeline, *R* genes were also identified in three other *Carya* species with assembled genomes: *C. illinoinensis*, *C. cathayensis*, and *C. sinensis*. In addition, we focused particularly on resistance to *Phylloxera* – aphid-like insects that induce gall formation in pecan. A major quantitative trait locus (QTL) associated with phylloxera resistance was identified by Lovell et al. (2021) in *C. illinoinensis*. Using the primary assembly of *C. illinoinensis* cv. ‘Lakota’ as the reference, this QTL is located on chromosome 16 (positions 1521681 to 2392040), between genes CiLak.16G012100 and CiLak.16G019000 (Lovell et al. 2021). Syntenic regions in *C. glabra* corresponding to this QTL were detected and visualized using MCScan from JCVI (v.1.2.10) (Tang et al. 2024). Within these syntenic regions, putative *R* genes were identified across all four *C. glabra* haplotypes.

## Results

### Statistics of sequence data

The basic statistics of the raw sequence data are summarized in Table 1. PacBio HiFi reads were generated on two SMRT cells, yielding a total of 79.1 gigabases (Gb) of data (44.1 Gb from one cell and 35.0 Gb from the other cell) (Table 1). In total, 5.3 million HiFi reads were obtained, with an average read length of 15.0 kilobases (kb). The proportions of bases with quality scores greater than 20 (Q20) and 30 (Q30) were 97.7% and 94.5%, respectively. The sequencing coverage, calculated by dividing the total number of bases by the monoploid genome size (1*x*), was 131.7× (Table 1). Given that the *Carya glabra* is a tetraploid and comprises four haplotypes, the coverage per haplotype was therefore 32.9×.

**Table 1.** Basic statistics of the raw sequence data from *Carya glabra*.

|  | PacBio HiFi | Omni-C | RNA-seq |  |
| --- | --- | --- | --- | --- |
|  |  |  | Leaf | Axillary bud |
| Total bases (Gb) | 79.1 | 39.7 | 24.3 | 22.5 |
| Total read number (million) | 5.3 | 264.5 | 160.6 | 148.8 |
| Average read length (bp) | 15,035.2 | 150.0 | 151.0 | 151.0 |
| Coverage* | 131.7× | 66.1× | - | - |

Note: \*sequencing coverage was calculated by dividing the total number of bases by the assembled monoploid (1x) genome size (600.4 Mb for haplotype A, as described in the nuclear genome assembly section).

For Omni-C data, a total of 264.5 million reads (derived from paired-end sequencing of 132.3 million DNA fragments) were generated, and the total number of bases was 39.7 Gb (Table 1). The Q20 and Q30 quality scores were 98.7% and 96.4%, respectively. Sequencing coverage was 66.1×, corresponding to 16.5× per haplotype in the tetraploid genome.

RNAs extracted from leaf and axillary bud tissues were of high quality, with RNA Integrity (RIN) scores of 7.1 and 7.2, respectively. For RNA-seq data, 161.6 million reads (from paired-end sequencing of 80.3 million fragments) were generated from the leaf tissue, and the Q20 and Q30 quality scores were 99.0% and 96.1%, respectively (Table 1). We also generated 148.8 million reads from the axillary bud tissue, and the Q20 and Q30 scores were 99.0% and 96.0%, respectively.

### Chloroplast and mitochondrial genome assembly and annotation

The chloroplast genome of *Carya glabra* is 160,839 bp in length and has the typical quadripartite structure (Fig. 2). The genome is composed of a pair of inverted repeat (IR) regions (i.e., IRA and IRB; 26,006 bp in length for each region), a large single-copy (LSC) region (90,041 bp), and a small single-copy (SSC) region (18,786 bp) (Fig. 2). A total of 113 unique genes, including 79 protein-coding genes, 30 tRNA genes, and 4 rRNA genes, were annotated (Fig. 2). A detailed list of these genes, along with their functional categories and genomic locations, is provided in Table S3. The GC contents of LSC, SSC, and IR regions were 33.7%, 29.9%, and 42.6%, respectively.

**Fig. 2.**
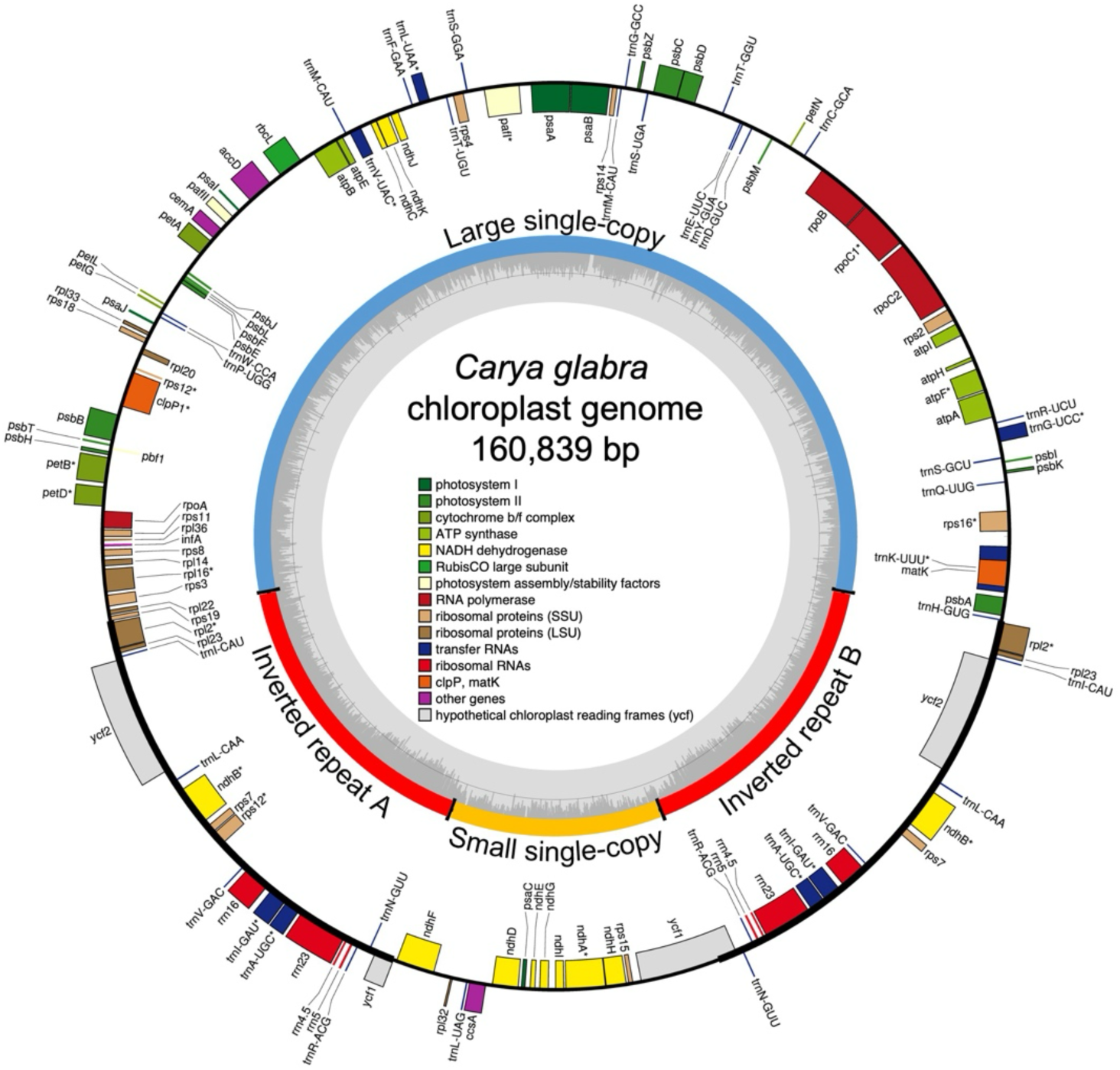
Annotated chloroplast genome of *Carya glabra*. The outermost circle shows the annotated genes, color-coded according to their functional categories (legend displayed in the figure center). Genes on the inside of the circle are transcribed clockwise, whereas those on the outside are transcribed counterclockwise. Intron-containing genes are marked with an asterisk (*). The inner circle indicates the four structural regions of the chloroplast genome: the large single-copy, the small single-copy, and the two inverted repeat regions (A and B). The innermost grey graph represents the GC content, with the grey reference line marking the 50% threshold. The figure is modified from the OGDRAW output.

The two mitochondrial chromosomes are 493,063 bp and 147,309 bp in length (Fig. 3). The larger chromosome (mtChr1) also presents a quadripartite structure where two inverted repeats (mtIR) of 2,760 bp intercalate a small single-copy (mtSSC) region (135,915 bp) and a large single-copy (mtLSC) region (351,628 bp). The smaller chromosome (mtChr2) is mostly redundant with mtChr1, consisting of one of the mtIRs, the entire mtSSC, 1,795 bp of the mtLSC, and a unique segment of 6,839 bp. A total of 42 protein-coding genes, 23 tRNA genes, and 3 rRNA genes were annotated in the mitochondrial genome (Table S4). From these, 15 were annotated as functional plastome-derived genes (5 protein-coding genes and 10 tRNA genes) (Table S4). We additionally identified 15 nonfunctional plastome genes: six were complete but contained premature stop codons, and nine were only fragmentary. All plastome-derived genes were located inside several sequence segments with varying lengths and degrees of conservation, as measured by their sequence identity to the chloroplast assembly (Table S5). Most notably, two large segments contained multiple functional plastome genes, the first segment (15,031 bp, 99.2% identity) contained *trnA-UGC*, *trnI-CAU*, *trnL-CAA*, and *trnV-GAC* genes; and the second segment (2,137 bp, 84.3% identity) contained *psaJ*, *rpl20*, and *rpl33* genes (Table S5).

**Fig. 3.**
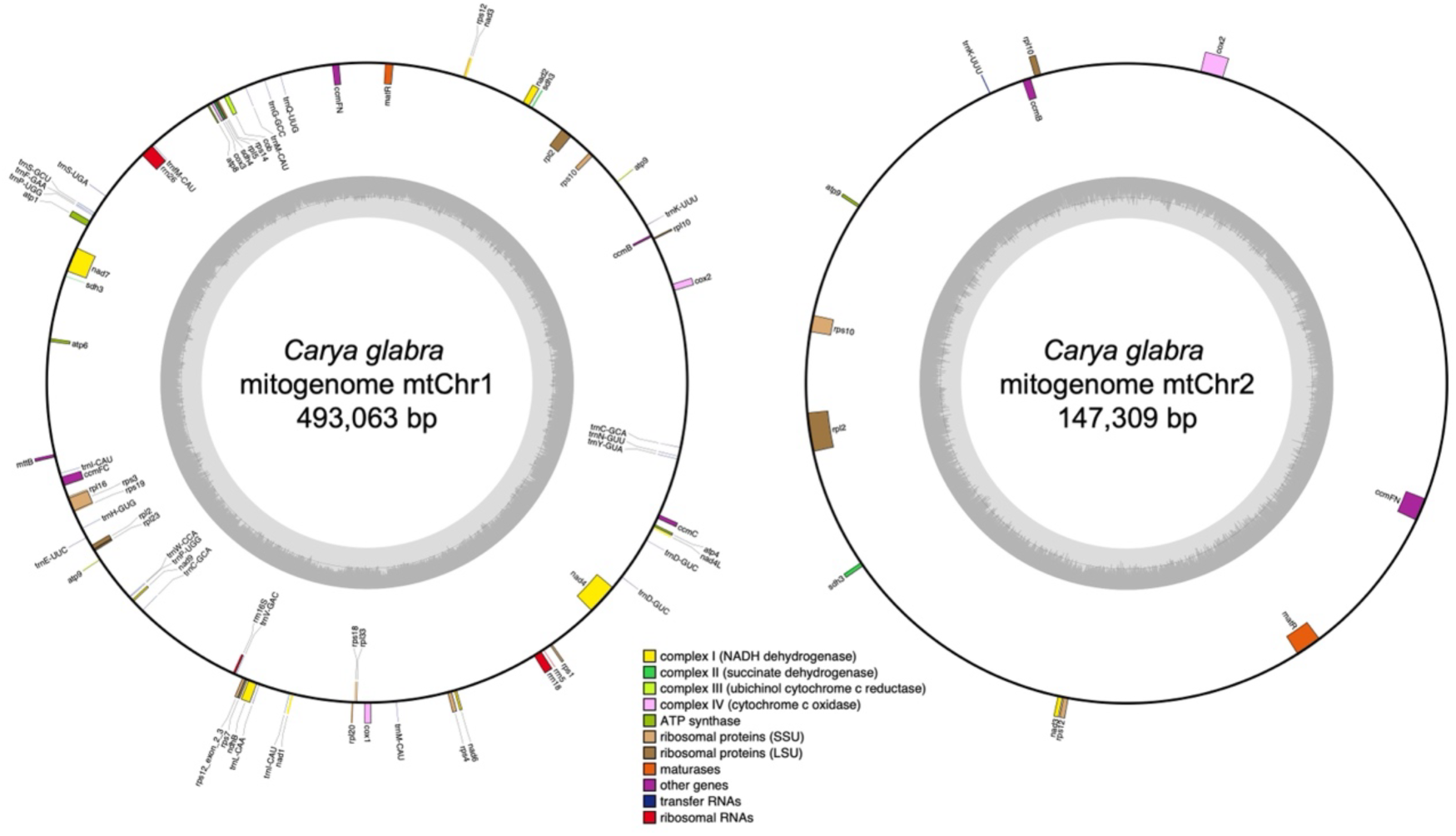
Annotated mitochondrial genome of *Carya glabra* shown as two conformations labeled mtChr1 and mtChr2. The outermost circle shows the annotated genes, color-coded according to their functional categories (legend displayed at bottom center). Genes on the inside of the circle are transcribed clockwise, whereas those on the outside are transcribed counterclockwise. The innermost grey graph represents the GC content, with the grey reference line marking the 50% threshold. Chromosomes are not drawn to scale. The figure is modified from the OGDRAW output.

### Nuclear genome profiling

Based on *k*-mer frequency analysis of the unassembled HiFi reads, GenomeScope 2.0 estimated the monoploid genome size as 515.4 Mb, with a heterozygosity value of 4.9% and repetitive sequences accounting for 38.5% of the genome. The frequencies of the heterozygous forms *aaab* and *aabb* were 3.2% and 1.4%, respectively. The resulting *k*-mer spectrum is shown in Fig. 4. The four major peaks, corresponding to *k*-mers present in one to four copies, are characteristic of an autotetraploid genome.

**Fig. 4.**
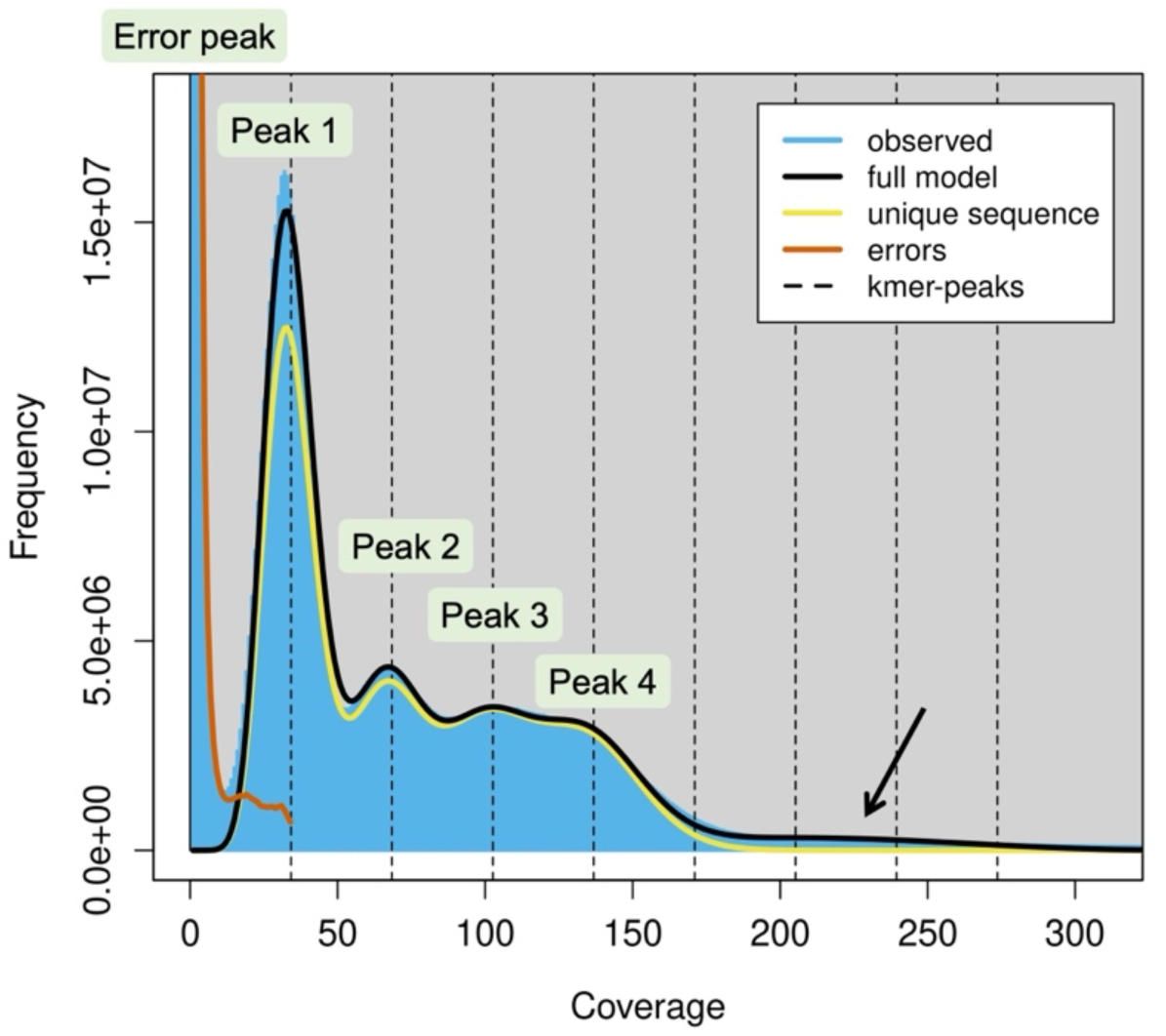
*K*-mer spectrum of *Carya glabra*. The plot illustrates the distribution of *k*-mer frequences (i.e., counts of unique *k*-mers; *y*-axis) across different coverage depths (*x*-axis) in the entire HiFi dataset. The leftmost error peak, representing the large number of low-coverage unique *k*-mers, results from sequencing errors. Peaks 1, 2, 3, and 4 correspond to *k*-mers present in one, two, three, and four copies, respectively, within the tetraploid genome. The coverages for peaks 1, 2, 3, and 4 are 34.2×, 68.4×, 102.6×, and 136.8×, respectively. The high-coverage “hump”, indicated by the arrow, represents *k*-mers derived from repetitive regions. *K*-mer size: 21. The figure is modified from the GenomeScope 2.0 output.

### Nuclear genome assembly and annotation

The initial unitig assembly generated by hifiasm comprised 2,856 unitigs with an N50 of 7.5 Mb. A dot plot comparing this unitig assembly with one set of chromosomes from the *Carya illinoinensis* genome revealed that each *C. illinoinensis* region corresponded to four unitigs, confirming the tetraploid nature of the *C. glabra* genome and indicating that the unitig assembly incorporated genomic sequences from all four haplotypes (Fig. S1). The complete BUSCO score for the unitig assembly was 98.9%, consisting of 1.0% single-copy and 97.9% duplicated BUSCOs; the high proportion of complete and duplicated BUSCOs reflects that sequences from all haplotypes were represented in the assembly.

Next, the unitigs were scaffolded by YaHS using the Omni-C data. Based on the hic_qc analysis, the Omni-C library was considered “sufficient”, showing high proportions of long-distance and inter-unitig contacts (Table S6). The initial YaHS scaffolding resulted in 2,584 scaffolds with an N50 of 36.9 Mb, including 62 scaffolds longer than 10 Mb. Examination of the Hi-C contact map, along with the dot plot comparing the *Carya illinoinensis* genome with the initial YaHS scaffolds, revealed several scaffolding errors, including two misjoins and an inversion error, which were corrected manually using Juicebox (Fig. S2). In addition, Juicebox was used to reorient several scaffolds to match the chromosome orientations of *C. illinoinensis*.

After manual curation, the final assembly contained 64 scaffolds longer than 10 Mb, accounting for 94.8% of the total assembled sequences (2,319.4 Mb out of 2,445.8 Mb) and corresponding to the expected chromosome number of the *Carya glabra* genome (Fig. 5). Hereafter, we refer to these 64 scaffolds as pseudo-chromosomes (or simply chromosomes for brevity). Each pseudo-chromosome was named according to its syntenic similarity with the *C. illinoinensis* genome based on the dot plot (Fig. 5c) and was assigned to haplotypes (A through D) based on descending length. It is important to note that this haplotype assignment is artificial and does not necessarily reflect true biological haplotypes (see Materials and Methods). The monoploid genome (1*x*) sizes for haplotypes A, B, C, and D were 600.4 Mb, 585.2 Mb, 574.3 Mb, and 559.4 Mb, respectively (Table 2). In addition, the complete BUSCO scores for the assembled genomes were 97.8%, 97.6%, 96.8%, and 95.4% for haplotypes A, B, C, and D, respectively (Table 2). Detailed statistics for each chromosome are provided in Table S7.

**Fig. 5.**
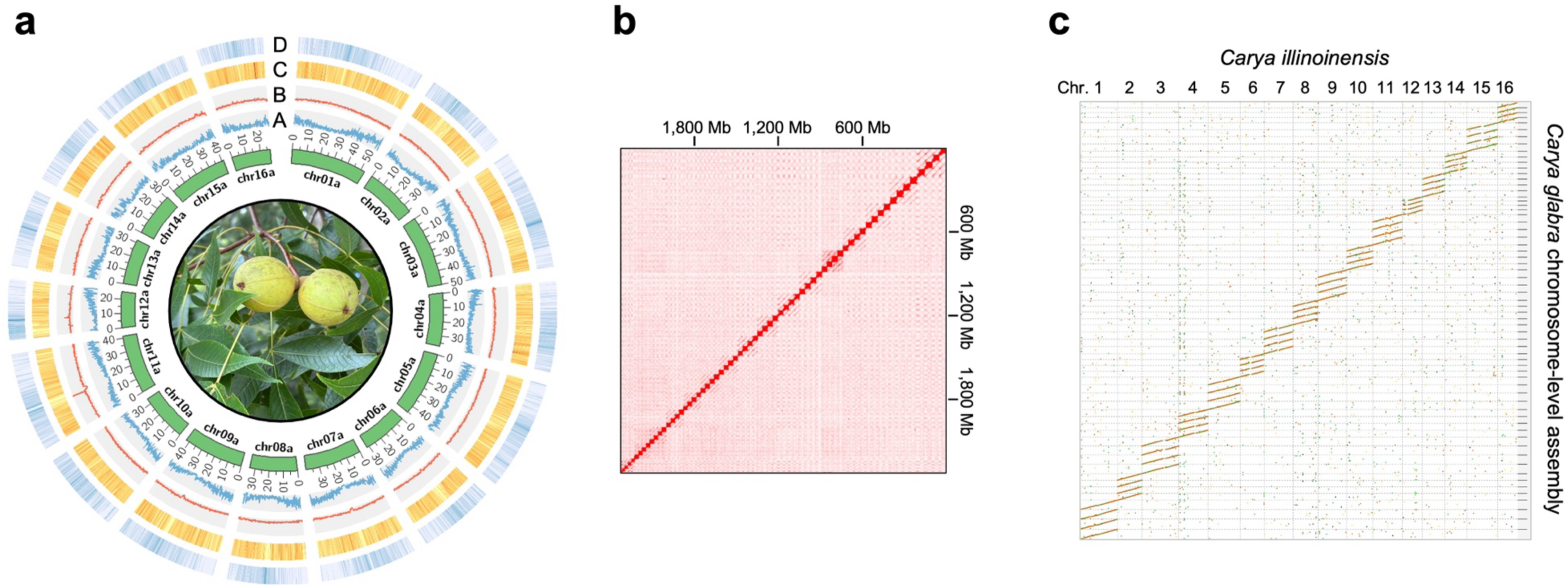
The chromosome-level assembly of the *Carya glabra* (4*x*) nuclear genome. (a) Circos plot of the 16 chromosomes from haplotype A of the *Carya glabra* genome. The unit of the chromosome length is Mb. The densities of various genomic features in 100-kb sliding windows across the chromosomes are shown on four tracks (A: genes; B: transposons; C: *copia*; D: *gypsy*). (b) The Hi-C contact map of the nuclear genome assembly. (c) The dot plot comparing one set of chromosomes from *Carya illinoinensis* (2*x*) and the four sets of chromosomes from *C. glabra*.

**Table 2.** Assembly statistics and genomic features of the *Carya glabra* genome and other published genomes of *Carya* species.

| Genome statistics | *C. glabra* (4x) |  |  |  | *C. illinoensis* (2x)<sup>1</sup> | *C. sinensis* (2x) | *C. cathayensis* (2x) |
| --- | --- | --- | --- | --- | --- | --- | --- |
|  | Hap. A | Hap. B | Hap. C | Hap. D |  |  |  |
| Monoploid (1x) genome size (Mb) | 600.4 | 585.2 | 574.3 | 559.4 | 674.3 | 623.2 | 698.1 |
| N50 (Mb) | 39.6 | 37.7 | 36.8 | 36.2 | 44.7 | 38.9 | 43.5 |
| Repeat sequences (%) | 55.0 | 54.4 | 54.0 | 53.8 | 49.7 | 43.5 | 50.1 |
| Predicted protein-coding genes | 30,947 | 31,087 | 30,369 | 30,110 | 32,267 | 35,370 | 36,722 |
| Complete BUSCO (%) assembly | 97.8 | 97.6 | 96.8 | 95.4 | 98.1 | 96.9 | 97.0 |
| Complete BUSCO (%) annotation | 97.7 | 97.1 | 96.5 | 94.9 | 96.3 | 94.8 | 95.8 |
| Reference | Current work |  |  |  | Lovell et al. 2021 | Zhang et al. 2024b |  |

Note: <sup>1</sup>the statistics are from *C. illinoensis* cv. ‘Pawnee’. Hap.: haplotype.

Repetitive sequences accounted for the majority of the *Carya glabra* genome (Table 2; Table S8). In haplotypes A, B, C, and D, 55.0%, 54.4%, 54.0%, and 53.8% of the genomic sequences were classified as repetitive regions, respectively (Table 2). Specifically, retrotransposons comprised 24.7-27.2% of the genome across the four haplotypes, and DNA transposons represented 19.4-21.5% of the genome (Table S8). In addition, simple repeats (duplications of short DNA motifs; microsatellites) accounted for 1.2-1.3% of the genome.

For protein-coding gene prediction, several BRAKER3 settings were tested using the haplotype A genome as the reference (Table S2). The combination that used RNA-seq data from *C. glabra* and protein evidence from 14 model species – followed by filtering out gene models ≤50 amino acids – produced the highest BUSCO score (97.7%) (Table S2). Therefore, the same setting was used to annotate the genes from haplotypes B, C, and D.

A total of 30,947 genes were predicted for haplotype A, with an average CDS length of 1,241 bp (Table 2; Table S9). For haplotypes B, C, and D, the number of predicted protein-coding genes ranged from 30,110 to 31,087 (Table 2). The average CDS length ranged from 1,239 bp to 1,254 bp (Table S9). All haplotypes had an average of 5.0 exons per gene, and the average gene length varied between 4,364 bp and 4,460 bp (Table S9).

TRAPID annotation assigned gene family information to 94.3% of the predicted genes in haplotype A, with 79.7% and 86.3% of genes annotated with Gene Ontology (GO) terms and protein domains, respectively (Table S9). The core gene family completeness score in TRAPID was 0.982, exceeding the conservation threshold of 0.9, further supporting the high completeness of the predicted gene models. Similarly, haplotypes B, C, and D showed high annotation rates: 93.9–94.7% of genes were assigned to gene families, and 85.9–86.5% were annotated with protein domains (Table S9). All haplotypes also exhibited high BUSCO completeness scores based on the annotated genes, ranging from 94.9% to 97.7% (Table 2).

### Comparative genomic analysis

Synteny analysis was performed among the four haplotypes of *Carya glabra* and the haploid genomes of *C. cathayensis*, *C. illinoinensis*, and *C. sinensis*, revealing high overall collinearity among the genomes (Fig. 6). However, several structural variants were also identified. For example, an inversion on chromosome 16 was detected between the *C. sinensis* and *C. illinoinensis* genomes (indicated by green circle 1 in Fig. 6). Another inversion on chromosome 11 was observed between *C. illinoinensis* and all four haplotypes of *C. glabra* (green circle 2); this inversion was also evident in the corresponding dot plot (Fig. 5c). Furthermore, structural variation was found among the four *C. glabra* haplotypes. For instance, between haplotypes B and C, the synteny analysis showed an inversion on chromosome 3, which was also detected in the dot plot (Fig. 5c; green circle 3 in Fig. 6).

**Fig. 6.**
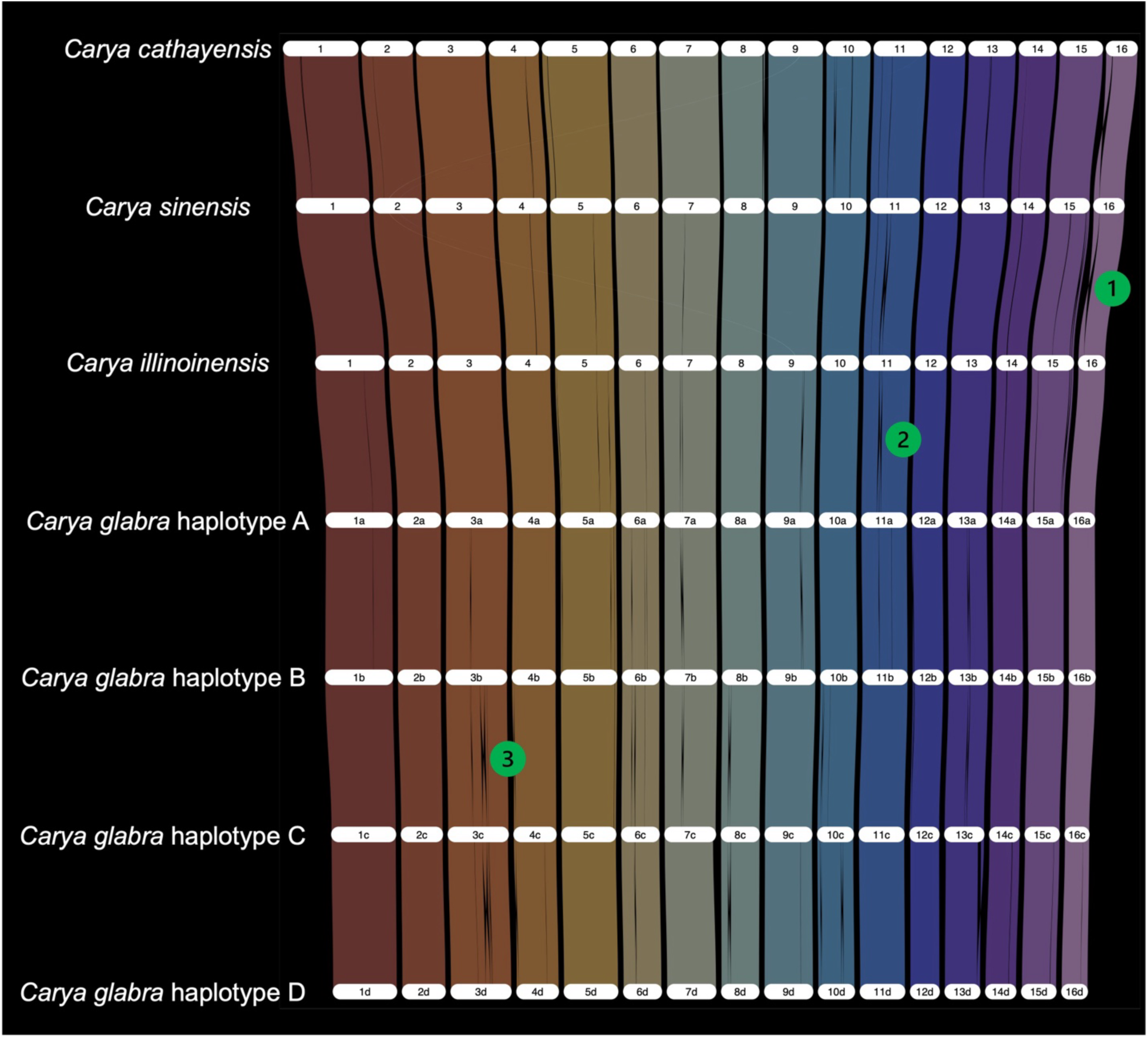
Syntenic map (riparian plot) of homologous regions among the four haplotypes of *Carya glabra* and the haploid genomes of *C. cathayensis*, *C. sinensis*, and *C. illinoinensis*. The chromosomes are scaled by gene rank order. Among the structural variants identified, three are highlighted: green circle 1 marks an inversion on chromosome 16 between *C. sinensis* and *C. illinoinensis*; green circle 2 indicates an inversion between *C. illinoinensis* and haplotype A of *C. glabra* on chromosome 11; an inversion between *C. glabra* haplotypes B and C on chromosome 3 is indicated by green circle 3.

### Disease resistance genes in *C. glabra*

Plant disease resistance genes, i.e., *R* genes, across the four haplotypes were predicted. Specifically, we focused on four major classes of *R* genes: CNL [containing the coiled-coil domain, the nucleotide-binding site (NBS) domain, and the leucine-rich repeat (LRR) domain], TNL (containing the Toll-interleukin receptor-like domain, the NBS domain, and the LRR domain), RLP [receptor-like protein, containing the transmembrane (TM) domain and the LRR domain], and RLK (receptor-like kinase, containing the TM domain, the LRR domain, and the kinase domain). In haplotype A, we identified 625 putative *R* genes from these four classes, including 56 CNL, 39 TNL, 214 RLP, and 316 RLK class genes (Table S10). For haplotypes B, C, and D, 638, 655, and 608 putative *R* genes were annotated, respectively (Table S10). In addition, we identified 724, 685, and 800 putative *R* genes in the primary assemblies of *C. illinoinensis*, *C. sinensis*, and *C. cathayensis*, respectively (Table S10).

The syntenic regions in *C. glabra* corresponding to the major QTL for phylloxera resistance in *C. illinoinensis* were identified on chromosome 16 (Fig. 7). Within these syntenic regions, 8, 10, 11, and 8 *R* genes were detected in haplotypes A, B, C, and D, respectively (Fig. 7; Table S11). Syntenic gene pairs between the five *R* genes annotated in the primary assembly of *C. illinoinensis* cv. ‘Lakota’ and their counterparts in *C. glabra* were highlighted in the synteny plot (Fig. 7). Among the 37 *C. glabra R* genes (30 of 37) located in these syntenic regions, 30 belong to the TNL class, while 3 and 4 belong to the RLP and RLK classes, respectively (Table S11).

**Fig. 7.**
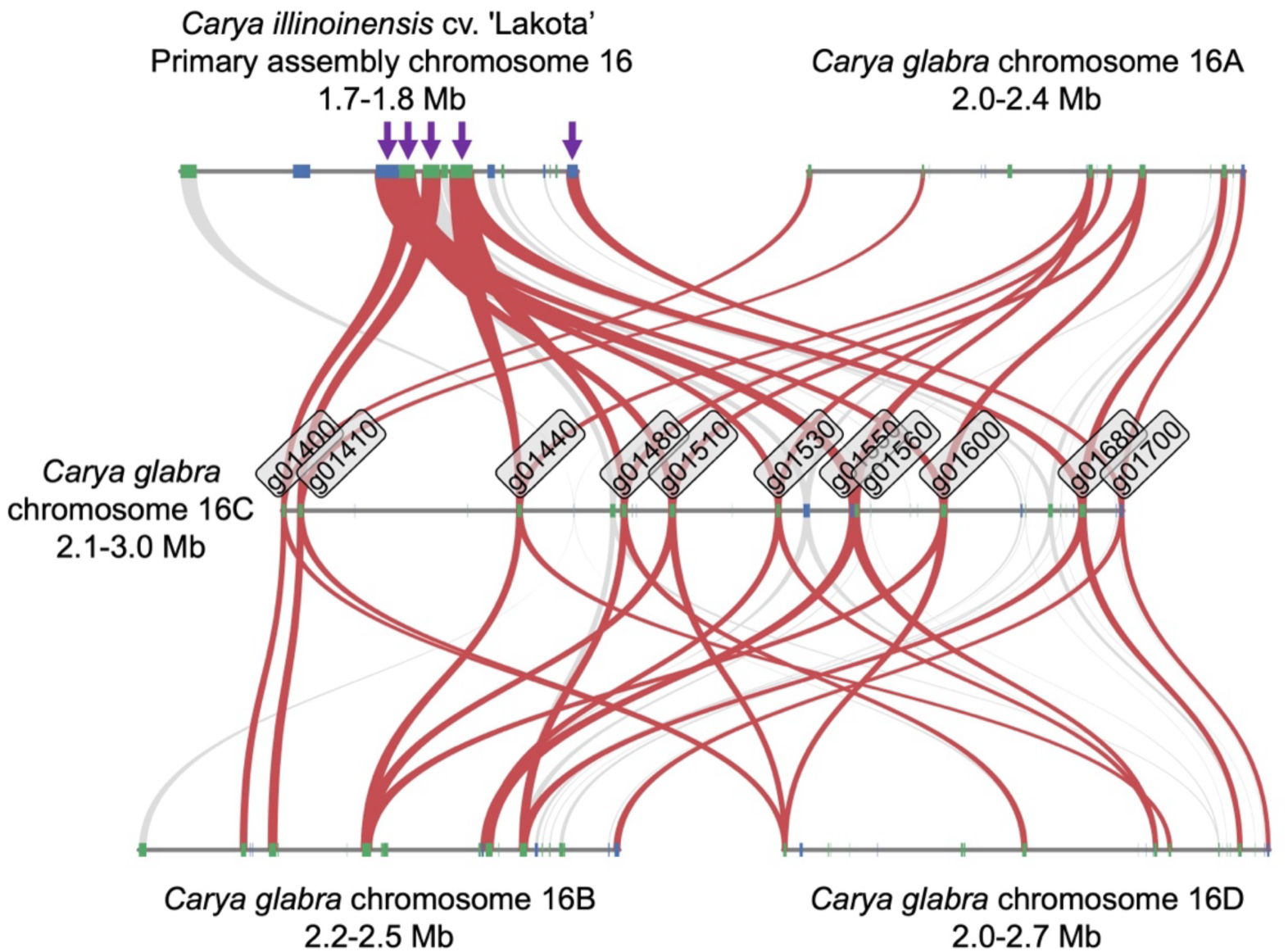
Synteny between *Carya illinoinensis* cv. ‘Lakota’ and the four *Carya glabra* haplotypes at the major quantitative trait locus (QTL) associated with phylloxera resistance. QTL mapping in *C. illinoinensis* by Lovell et al. (2021) identified a single large QTL peak on chromosome 16. Within this QTL on the primary assembly of *C. illinoinensis* cv. ‘Lakota’, five putative plant disease resistance genes (*R* genes) containing the leucine-rich repeat (LRR) domain were annotated (indicated by arrowheads). In the corresponding syntenic region of *C. glabra*, chromosome 16C contains 11 putative *R* genes – the highest count among the four haplotypes – with each gene labeled by name. The syntenic regions on chromosomes 16A, 16B, and 16D contain 8, 10, and 8 *R* genes, respectively. Syntenic gene pairs are connected by the ribbons, with those linking to the 11 *R* genes on *C. glabra* chromosome 16C highlighted in red. Note that not all *R* genes on chromosomes 16A, 16B, and 16D are reciprocal best hits with *R* genes on chromosome 16C; therefore, these are not connected with red ribbons in the plot. Genes are depicted as boxes, with blue representing genes on the positive strand and green representing genes on the negative strand. Chromosome segments are not drawn to scale.

## Discussion

### *Carya glabra* organellar genomes

The chloroplast genome size in Juglandaceae ranges from 158,223 bp to 161,713 bp (Liu et al. 2025). Three *Carya glabra* chloroplast genomes have been published to date (Luo et al. 2021; Xi et al. 2022; Liu et al. 2025), with sizes ranging from 160,645 bp to 160,652 bp. In the present study, the assembled chloroplast genome of *C. glabra* is 160,839 bp in length (Fig. 2), very similar to the published *C. glabra* chloroplast genomes and within the size range observed across species from other Juglandaceae.

A total of 109, 113, and 114 unique genes were annotated in previously published *C. glabra* chloroplast genomes with NCBI accession numbers OR099205, NC_067504, and BK061156, respectively. In our study, 113 unique genes were identified, including 79 protein-coding genes, 30 tRNA genes, and 4 rRNA genes (Fig. 2; Table S3). The additional gene reported in accession BK061156 is *ycf15*, a functionally uncharacterized gene that is also absent from the well-annotated *Nicotiana tabacum* chloroplast genome (NC_001879). Through manual curation, we identified several misannotated and missing genes in previously reported *C. glabra* chloroplast genomes (summarized in Table S12). For example, additional copies of tRNA genes *trnA-UGC* and *tnrM-CAU* were misannotated in BK061156; two protein-coding genes, *atpB* and *rpoB*, were missing from OR099205; and the first exons of *petB*, *petD*, and *rpl16* were absent from NC_067504. All such potential annotation errors were manually corrected in the present study. Together, these results indicate that although several *C. glabra* chloroplast genomes have been published, our assembly and annotation represent the most complete and accurate version to date.

Compared to chloroplast genomes, the reports of the assembly of plant mitochondrial genomes are few, primarily due to the high structural complexity of the mitogenome in plants (Palmer and Herbon 1988; Møller et al. 2021; Wu et al. 2022; Wang et al. 2024). Only a few mitochondrial genomes have been published for species from Juglandaceae, and those available mitogenomes show substantial variation in structure and gene content. Chen et al. (2024) assembled the first mitochondrial genome of *Carya illinoinensis*: the single circular genome is 495.2 kb in length and contains 37 protein-coding genes, 24 tRNA genes, and 3 rRNA genes. The *Juglans regia* (Juglandaceae) mitogenome consists of three circular chromosomes and includes 39 protein-coding genes, 47 tRNA genes, and 5 rRNA genes (Ye et al. 2024). The *Juglans mandshurica* mitochondrial genome includes two chromosomes and has 38 protein-coding genes, 20 tRNA genes, and 3 rRNA genes (Su et al. 2023). In *Carya glabra*, the mitogenome includes two chromosomes (493.1 kb and 147.3 kb in length), and we identified 42 protein-coding genes, 23 tRNA genes, and 3 rRNA genes (Fig. 3; Table S4). Although mitogenomes are generally highly variable, the *C. glabra* mitochondrial genome is broadly comparable with other published Juglandaceae mitogenomes.

The varying sizes and identities of the plastome segments detected in the *C. glabra* mitochondrial genome suggest multiple transfer events occurring at different times (Table S5). In future studies, it would be interesting to compare these transferred segments with other congeneric chloroplast and mitochondrial genomes.

### Nuclear genomes in *Carya*

We assembled and annotated the first nuclear genome of *Carya glabra* (Fig. 5). The assembly is chromosome-level and haplotype-resolved, representing the first assembled polyploid genome in the genus (Fig. 5). Furthermore, GenomeScope 2.0 predicted that *Carya glabra* is an autotetraploid based on the pattern of nucleotide heterozygosity levels: the frequency of the heterozygous *aaab* genotype was higher than that of the *aabb* genotype (3.2% versus 1.4%), a pattern characteristic of autopolyploids (Ranallo-Benavidez et al. 2020). Additionally, the *k*-mer spectrum showing four major peaks (Fig. 4), along with the high similarity among the four copies of each chromosome compared to the *C. illinoinensis* genome based on the dot plot (Fig. 5c), further support that *C. glabra* is an autotetraploid.

In terms of genomic composition, 53.8-55.0% of the *Carya glabra* genome consists of repetitive sequences, with slight variation among haplotypes (Table 2). Similar, but lower, proportions of repetitive content have been reported in other *Carya* species. Lovell et al. (2021) found that 49.7% of the *C. illinoinensis* genome is repetitive sequences, and Zhang et al. (2024b) reported repeat fractions in the genomes of *C. sinensis* (43.5%) and *C. cathayensis* (50.1%) (Table 2).

We predicted more than 30,000 protein-coding genes for each *Carya glabra* haplotype (Table 2). BUSCO completeness scores were high across all haplotypes, with haplotype A having a BUSCO score of 97.7%. The number of genes predicted in *Carya glabra* is broadly comparable to those reported for other *Carya* species (Table 2). Lovell et al. (2021) annotated 32,267 genes in *C. illinoinensis*, and Zhang et al. (2024b) identified 35,370 and 36,722 genes in *C. sinensis* and *C. cathayensis*, respectively (Zhang et al. 2024b).

Several non-mutually exclusive factors may explain the differences in gene count among *Carya* genomes. First, the annotation pipeline can affect the number of predicted genes. Weisman et al. (2022) found that applying different annotation methods to the same genome can lead to the identification of genes unique to each method. In this study, we used BRAKER3 for gene annotation, whereas PASA (Haas et al. 2003) and FGENESH (Salamov et al. 2020) were used to annotate the *C. illinoinensis* genome (Lovell et al. 2021). Zhang et al. (2024b) used PASA, AUGUSTUS (Stanke et al. 2006), and GeneWise (Birney et al. 2004) to annotate the *C. sinensis* and *C. cathayensis* genomes. Second, the diversity and number of tissues represented in the RNA-seq data can affect annotation completeness, and sampling from multiple tissues is recommended (Salzberg 2019; Kress et al. 2022; Vuruputoor et al. 2023). Our annotations were supported by RNA-seq data from two tissues (leaf and axillary bud), whereas Lovell et al. (2021) used RNA-seq data from a larger number of tissues, including leaf, catkin, and dormant and swelling buds. Lastly, the lower gene count in *C. glabra* may reflect its polyploid nature. Genome fractionation and gene loss are common following polyploid formation (Langham et al. 2004; Leitch and Bennett 2004; Freeling 2009; Soltis et al. 2015; Van de Peer et al. 2017; Wendel et al. 2018), although fractionation as originally defined (Freeling 2009) cannot occur in an autopolyploid that lacks parental subgenomes. Indeed, the relatively smaller monoploid (1*x*) genome size of *C. glabra* (e.g., 600.4 Mb for haplotype A and smaller for the other haplotypes) compared with diploid *Carya* species (e.g., 674.3 Mb for *C. illinoinensis*) may result from gene loss following polyploidy in *C. glabra*.

In summary, the *C. glabra* genome assembly and annotation presented in this study are of high quality, with metrics comparable to, or surpassing (based on the BUSCO completeness score; Table 2), published genomes from other *Carya* species.

### Potential practical applications of the *Carya glabra* genome assembly

The *Carya glabra* genome assembly provides a valuable resource for identifying candidate genes that may facilitate breeding programs in pecan (*C. illinoinensis*) and Chinese hickory (*C. cathayensis*). Notably, we identified over 600 disease resistance genes (*R* genes) in each haplotype of *C. glabra* (Table S10). A similar, but higher, number of *R* genes has been identified in other *Carya* species: *C. illinoinensis*, *C. sinensis*, and *C. cathayensis* have 724, 685, and 800 *R* genes, respectively (Table S10). We focused particularly on a genomic region syntenic to a major QTL associated with phylloxera resistance in *C. illinoinensis*. Several aphid-like insects from the genus *Phylloxera* infect pecan and induce gall formation, which can cause defoliation and significantly reduce yield (Hedin et al. 1985; Andersen and Mizell III 1987). Lovell et al. (2021) identified a single major QTL underlying this trait, and several candidate *R* genes containing LRR domains were annotated within this QTL. In the syntenic region in *C. glabra*, we identified 8, 10, 11, and 8 *R* genes in haplotypes A, B, C, and D, respectively (Fig. 7; Table S11). These candidate genes provide an additional genetic resource that could facilitate engineering efforts to improve phylloxera resistance in pecan.

Polyploidy plays an important role in plant breeding (Udall and Wendel 2006; Sattler et al. 2016), and polyploids often exhibit an advantageous stress response relative to diploids (Bomblies 2020; Fox et al. 2020; Van de Peer et al. 2021; Tossi et al. 2022). Future studies examining stress response in *Carya glabra* and its closely related diploid species (e.g., *C. palmeri* and *C. illinoinensis*) could provide valuable insights into the effect of polyploidy on stress tolerance in *Carya* – information that may inform future strategies for improving pecan and Chinese hickory.

### Genome assembly and annotation as tools for conservation and teaching genomics

McCarty Woods is a 2.9-acre (11,735.9 m^2^) designated Conservation Area located at the heart of the UF campus (Fig. 1b). Representing part of the southernmost extent of deciduous forest in eastern North America, McCarty Woods contains more than 100 native plant species, including *Carya glabra* (Sharman 2024). Although designated as a Conservation Area, McCarty Woods’ central location on the UF campus has made it a recurring target for development. In 2021, a campaign led by botanists at the Florida Museum of Natural History as well as students and community members successfully halted proposed development plans, and efforts to advocate for long-term protection and restoration of the Woods are ongoing.

In collaboration with the ACTG project, the McCarty Woods Genome Project launched in 2024 (Sharman 2024). By sequencing the first genomes of iconic trees growing in the Woods, the project aims to “immortalize” these individuals and provide reference genomes that will guide future research and applications involving these species. These genomic resources strengthen the case for preserving the Conservation Area status for McCarty Woods and underscore its significant value for research and education. The reference genome of *Carya glabra* presented in this study represents the first genome produced by the McCarty Woods Genome Project, with others in progress (e.g., *Quercus michauxii*).

A Course-based Undergraduate Research Experience (CURE) class was offered at UF in Spring 2025 as part of the McCarty Woods Genome Project (Fig. 1c). Teaching materials and data analysis pipelines from the ACTG project (Harkess 2022; Yocca et al. 2024; Zhang et al. 2024a) were incorporated into the course, providing undergraduate students with hands-on experience in genome assembly and annotation of *Carya glabra*. By combining real-world data with active learning, the course engaged students from eight departments — Biology, Biomedical Engineering, Chemistry, Computer & Information Science & Engineering, English, Entomology and Nematology, Mechanical and Aerospace Engineering, and Statistics — and emphasized programming, collaboration, critical thinking, and scientific writing. Bioinformatic code generated through the course is publicly available on GitLab (https://gitlab.com/shengchenshan/bot4935-plant-genome-assembly-and-annotation), and lecture slides are available on Zenodo (https://doi.org/10.5281/zenodo.17969442). In summary, the course provided students insight into the process of scientific research and the role of genomics in biological sciences, highlighting the value of genome assembly and annotation in training the next generation of biological scientists and bioinformaticians.

## Future directions

The *Carya glabra* nuclear genome assembly provides an important tool for investigating the roles of polyploidy and hybridization in genome evolution in *Carya*. Several intriguing evolutionary questions remain. When did *C. glabra* undergo the most recent whole-genome duplication? Phylogenetic studies suggest that its closest relative is *C. texana*, which is also a tetraploid (Huang et al. 2019; Xi et al. 2022). Did these two species share an ancestral polyploidization event prior to divergence, or did they experience independent whole-genome duplication events? If the latter is the case, what is the diploid ancestor of *Carya glabra*? Are there undetected diploid populations of *C. glabra*? What environmental factors may have contributed to the success of genome doubling in these lineages?

The possibility of gene flow between *C. glabra* and pecan (*C. illinoinensis*), which is a diploid, also merits investigation. Plastome-based phylogenetic analyses have shown that *C. glabra* is closely related to a specific *C. illinoinensis* cultivar, ‘87MX3-2.11’ (Xi et al. 2022). If introgression involving *C. glabra* and pecan occurred, it may provide novel opportunities for pecan breeding and the potential transfer of beneficial traits from *C. glabra* into this economically important crop.

## Data availability

Raw data generated in this project, including PacBio HiFi, Omni-C, and RNA-seq, are deposited in NCBI under BioProject PRJNA1373287. The four haplotypes of the nuclear genome assembly are available under BioProject PRJNA1376128–PRJNA1376131. The nuclear genome annotation and organellar genomes are available at Zenodo (https://doi.org/10.5281/zenodo.17969322). All codes and scripts are available at: https://gitlab.com/shengchenshan/bot4935-plant-genome-assembly-and-annotation.

## Acknowledgments

The authors acknowledge Matthew A. Gitzendanner, Andre S. Chanderbali, and Lawrence Oshins from the University of Florida Research Computing team for their technical assistance and support. We also appreciate the helpful discussions with Rhett M. Rautsaw and Shujun Ou on PacBio sequencing and transposable element annotation, respectively. Computational resources were provided by HiPerGator, the University of Florida supercomputer.

## Author contributions

DES, PSS, AH, SS, and EMO designed the project. SS, EMO, PSS, DES, AH, BK, AO, GS, BS, RT, AT, EL, BP, TR, LS, GV, LW, and HZ contributed to data analysis and interpretation. SS, EMO, DES, PSS, HZ, AH, BK, AO, GS, BS, RT, AT, BP, TR, MHR, and GV wrote the manuscript. All authors reviewed and approved the manuscript.

## Supplementary materials

Fig. S1. Dot plot comparing one set of chromosomes from *Carya illinoinensis* (2*x*) with the unitig assembly of *Carya glabra* (4*x*).

Fig. S2. Manual curation of the YaHS scaffolding output using Juicebox.

Table S1. Protein evidence used for nuclear genome annotation.

Table S2. Statistics of gene models predicted under different BRAKER3 parameter settings for *Carya glabra* haplotype A.

Table S3. Annotated genes in the *Carya glabra* chloroplast genome.

Table S4. Annotated genes in the *Carya glabra* mitchondrial genome.

Table S5. Chloroplast-derived segments in the Carya glabra mitochondrial genome.

Table S6. Omni-C library quality control report from Phase Genomics’ hic_qc pipeline.

Table S7. Lengths (in Mb) of the 64 assembled pseudo-chromosomes of *Carya glabra*.

Table S8. Summary of repetitive element annotation in *Carya glabra*.

Table S9. Statistics of finalized gene models predicted for four haplotypes from *Carya glabra*.

Table S10. Four major classes of plant disease resistance genes (*R* genes) identified in *Carya glabra* and three other *Carya* species with assembled genomes.

Table S11. Putative *Carya glabra* plant disease resistance genes (*R* genes) identified in the syntenic regions corresponding to the major quantitative trait locus (QTL) associated with phylloxera resistance in *Carya illinoinensis*.

Table S12. Misannotated and missing genes in previously published *Carya glabra* chloroplast genomes.

