## Supplementary material for "A haplotype-resolved, chromosome-scale genome assembly and annotation for *Carya glabra* (pignut hickory; Juglandaceae)": Fig. S1

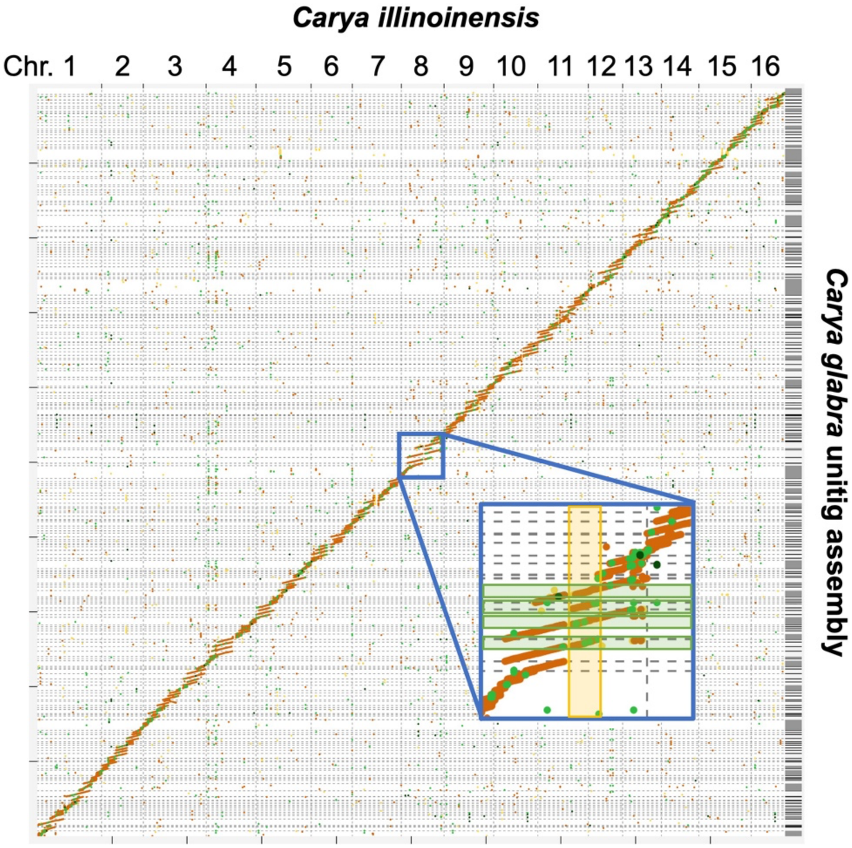


Fig. S1. Dot plot comparing one set of chromosomes from *Carya illinoinensis* (2*x*) with the unitig assembly of *Carya glabra* (4*x*). The zoomed-in panel shows that each *C. illinoinensis* region (shaded in yellow) corresponds to four unitigs from *C. glabra* (shaded in green), a pattern observed across the entire *C. illinoinensis* genome.
