## Supplementary material for "A haplotype-resolved, chromosome-scale genome assembly and annotation for *Carya glabra* (pignut hickory; Juglandaceae)": Fig. S2

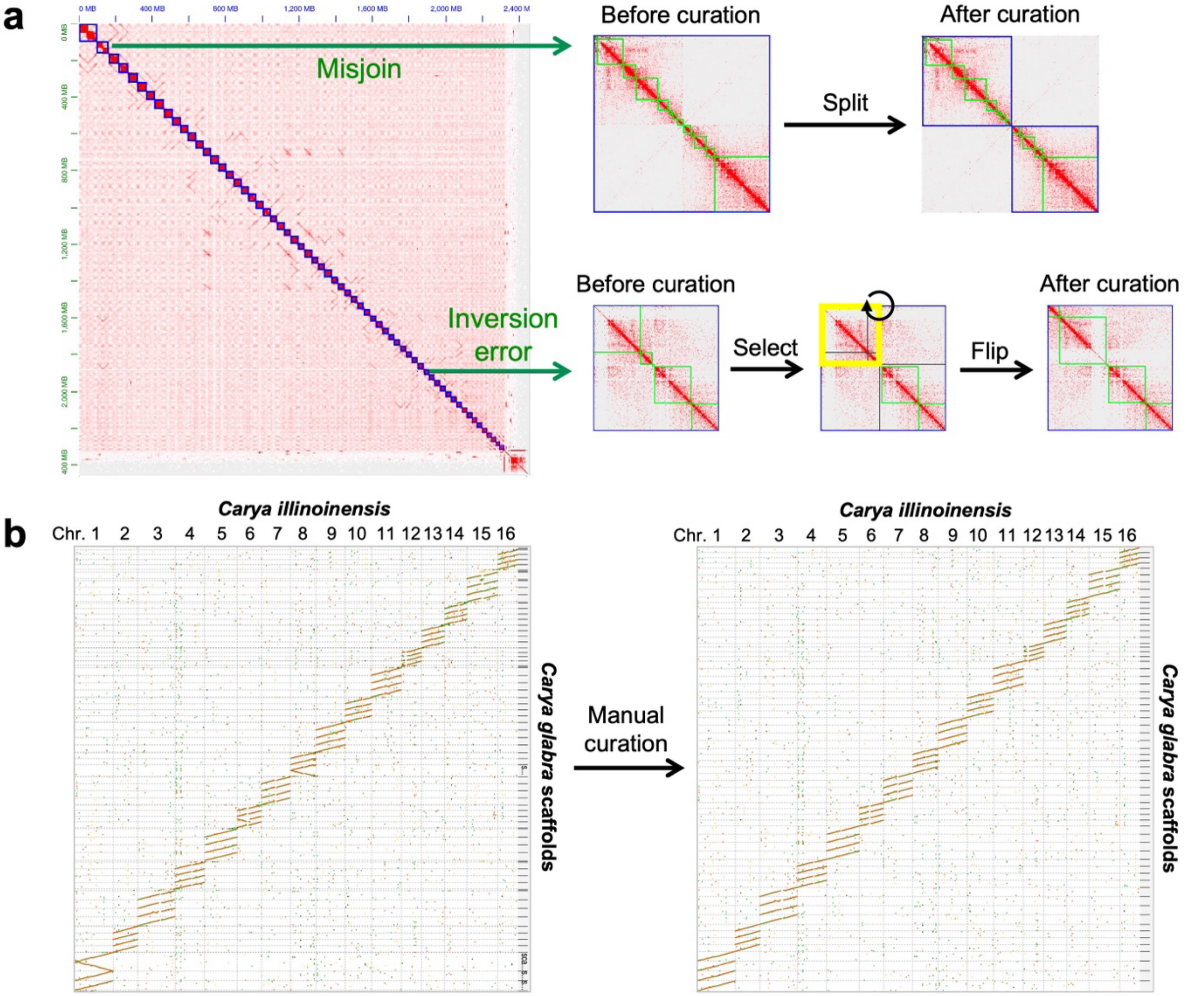


Fig. S2. Manual curation of the YaHS scaffolding output using Juicebox. (a) The Hi-C contact map generated in Juicebox was used to identify scaffolding errors. Two examples, one misjoin and one inversion error, are shown. In the contact map, each blue square represents a scaffold, and the green squares within a scaffold represent unitigs anchored to that scaffold. Based on the Hi-C signals, the misjoined scaffold was split into two scaffolds, while for the scaffold with an inversion error, the inverted unitigs (yellow square) were selected and inverted. (b) Dot plots comparing the *Carya illinoinensis* genome with the *Carya glabra* scaffolds before and after manual curation are shown.
