## Supplementary material for "A haplotype-resolved, chromosome-scale genome assembly and annotation for *Carya glabra* (pignut hickory; Juglandaceae)": Table S1

Table S1. Protein evidence used for nuclear genome annotation.

| Species | Source | Version/Accession |
|---|---|---|
| *Arabidopsis thaliana* (Brassicaceae) | Phytozome | Araport11 |
| *Cucumis sativus* (Cucurbitaceae) | Phytozome | v1.0 |
| *Eucalyptus grandis* (Myrtaceae) | Phytozome | v2.0 |
| *Glycine max* (Fabaceae) | Phytozome | Wm82.a6.v1 |
| *Malus domestica* (Rosaceae) | Phytozome | v1.1 |
| *Populus trichocarpa* (Salicaceae) | Phytozome | v4.1 |
| *Prunus persica* (Rosaceae) | Phytozome | v2.1 |
| *Vitis vinifera* (Vitaceae) | Phytozome | v2.1 |
| *Carya cathayensis* (Juglandaceae) | Juglandaceae Genome Projects[1] | v1.0 |
| *Carya illinoinensis* (Juglandaceae) | Phytozome | Pawnee v1.1 |
| *Carya sinensis* (Juglandaceae) | Juglandaceae Genome Projects | v1.0 |
| *Engelhardia roxburghiana* (Juglandaceae) | Juglandaceae Genome Projects | v1.0 |
| *Juglans regia* (Juglandaceae) | NCBI | GCA_001411555.2 |
| *Pterocarya stenoptera* (Juglandaceae) | Juglandaceae Genome Projects | v2.0 |

Note: [1]https://cmb.bnu.edu.cn/juglans/
