## Supplementary material for "A haplotype-resolved, chromosome-scale genome assembly and annotation for *Carya glabra* (pignut hickory; Juglandaceae)": Table S2

Table S2. Statistics of gene models predicted under different BRAKER3 parameter settings for *Carya glabra* haplotype A.

| Method | RNA-Seq data from *C. glabra* and protein data from nine model species[1] | | | RNA-Seq data from *C. glabra* and protein data from OrthoDB[2] (adding *C. illinoensis*) | | | RNA-Seq data from *C. glabra* and protein data from 11 model species[3] | | | RNA-Seq data from *C. glabra* and protein data from 14 model species[4] | | |
|---|---|---|---|---|---|---|---|---|---|---|---|---|
| Filter | None | 100 aa[5] | 50 aa[6] | None | 100 aa | 50 aa | None | 100 aa | 50 aa | None | 100 aa | 50 aa |
| Gene no. | 29,496 | 27,756 | 29,379 | 30,574 | 28,573 | 30,440 | 31,294 | 29,140 | 31,160 | 31,085 | 28,996 | 30,947 |
| Mean gene length (bp) | 4,558 | 4,787 | 4,574 | 4,444 | 4,698 | 4,461 | 4,357 | 4,620 | 4,374 | 4,380 | 4,639 | 4,398 |
| Mean CDS length (bp) | 1,268 | 1,333 | 1,273 | 1,248 | 1,319 | 1,253 | 1,229 | 1,302 | 1,234 | 1,236 | 1,308 | 1,241 |
| Average exons per gene | 5.2 | 5.4 | 5.2 | 5.0 | 5.3 | 5.0 | 5.0 | 5.2 | 5.0 | 5.0 | 5.2 | 5.0 |
| BUSCO | 97.6% | 97.1% | 97.6% | 97.5% | 97.1% | 97.5% | 97.6% | 97.1% | 97.6% | 97.7% | 97.1% | 97.7% |

Note: the statistics are based on the longest isoform of each gene. [1]the nine model species include *Arabidopsis thaliana*, *Cucumis sativus*, *Eucalyptus grandis*, *Glycine max*, *Malus domestica*, *Populus trichocarpa*, *Prunus persica*, *Vitis vinifera*, and *Carya illinoinensis*. [2]Viridiplantae database from the OrthoDB v.12 was used. [3]the 11 species include *Arabidopsis thaliana*, *Cucumis sativus*, *Eucalyptus grandis*, *Glycine max*, *Malus domestica*, *Populus trichocarpa*, *Prunus persica*, *Vitis vinifera*, *Carya cathayensis*, *Carya illinoinensis*, and *Carya sinensis*. [4]the 14 model species include *Arabidopsis thaliana*, *Cucumis sativus*, *Eucalyptus grandis*, *Glycine max*, *Malus domestica*, *Populus trichocarpa*, *Prunus persica*, *Vitis vinifera*, *Carya cathayensis*, *Carya illinoinensis*, *Carya sinensis*, *Engelhardia roxburghiana*, *Juglans regia*, and *Pterocarya stenoptera*. [5]gene models meeting any of the following criteria were filtered out: (1) presence of a premature stop codon; (2) absence of a start and/or stop codon; or (3) an open reading frame (ORF) length of ≤100 amino acids. [6]gene models meeting any of the following criteria were filtered out: (1) presence of a premature stop codon; (2) absence of a start and/or stop codon; or (3) an ORF length of ≤50 amino acids.
