## Supplementary material for "A haplotype-resolved, chromosome-scale genome assembly and annotation for *Carya glabra* (pignut hickory; Juglandaceae)": Table S3

Table S3. Annotated genes in the *Carya glabra* chloroplast genome. LSC: large single copy; SSC: small single copy; IR: inverted repeat.

| No. | Gene | Type | Location |
|---|---|---|---|
| 1 | *accD* | Protein coding | LSC |
| 2 | *atpA* | Protein coding | LSC |
| 3 | *atpB* | Protein coding | LSC |
| 4 | *atpE* | Protein coding | LSC |
| 5 | *atpF* | Protein coding | LSC |
| 6 | *atpH* | Protein coding | LSC |
| 7 | *atpI* | Protein coding | LSC |
| 8 | *ccsA* | Protein coding | SSC |
| 9 | *cemA* | Protein coding | LSC |
| 10 | *clpP1* | Protein coding | LSC |
| 11 | *infA* | Protein coding | LSC |
| 12 | *matK* | Protein coding | LSC |
| 13 | *ndhA* | Protein coding | SSC |
| 14 | *ndhB* | Protein coding | IR |
| 15 | *ndhC* | Protein coding | LSC |
| 16 | *ndhD* | Protein coding | SSC |
| 17 | *ndhE* | Protein coding | SSC |
| 18 | *ndhF* | Protein coding | SSC |
| 19 | *ndhG* | Protein coding | SSC |
| 20 | *ndhH* | Protein coding | SSC |
| 21 | *ndhI* | Protein coding | SSC |
| 22 | *ndhJ* | Protein coding | LSC |
| 23 | *ndhK* | Protein coding | LSC |
| 24 | *pafI* | Protein coding | LSC |
| 25 | *pafII* | Protein coding | LSC |
| 26 | *pbf1* | Protein coding | LSC |
| 27 | *petA* | Protein coding | LSC |
| 28 | *petB* | Protein coding | LSC |
| 29 | *petD* | Protein coding | LSC |
| 30 | *petG* | Protein coding | LSC |
| 31 | *petL* | Protein coding | LSC |
| 32 | *petN* | Protein coding | LSC |
| 33 | *psaA* | Protein coding | LSC |
| 34 | *psaB* | Protein coding | LSC |
| 35 | *psaC* | Protein coding | SSC |
| 36 | *psaI* | Protein coding | LSC |
| 37 | *psaJ* | Protein coding | LSC |
| 38 | *psbA* | Protein coding | LSC |
| 39 | *psbB* | Protein coding | LSC |
| 40 | *psbC* | Protein coding | LSC |
| 41 | *psbD* | Protein coding | LSC |
| 42 | *psbE* | Protein coding | LSC |
| 43 | *psbF* | Protein coding | LSC |
| 44 | *psbH* | Protein coding | LSC |
| 45 | *psbI* | Protein coding | LSC |
| 46 | *psbJ* | Protein coding | LSC |
| 47 | *psbK* | Protein coding | LSC |
| 48 | *psbL* | Protein coding | LSC |
| 49 | *psbM* | Protein coding | LSC |
| 50 | *psbT* | Protein coding | LSC |
| 51 | *psbZ* | Protein coding | LSC |
| 52 | *rbcL* | Protein coding | LSC |
| 53 | *rpl14* | Protein coding | LSC |
| 54 | *rpl16* | Protein coding | LSC |
| 55 | *rpl2* | Protein coding | IR |
| 56 | *rpl20* | Protein coding | LSC |
| 57 | *rpl22* | Protein coding | LSC |
| 58 | *rpl23* | Protein coding | IR |
| 59 | *rpl32* | Protein coding | SSC |
| 60 | *rpl33* | Protein coding | LSC |
| 61 | *rpl36* | Protein coding | LSC |
| 62 | *rpoA* | Protein coding | LSC |
| 63 | *rpoB* | Protein coding | LSC |
| 64 | *rpoC1* | Protein coding | LSC |
| 65 | *rpoC2* | Protein coding | LSC |
| 66 | *rps11* | Protein coding | LSC |
| 67 | *rps12* | Protein coding | IR |
| 68 | *rps14* | Protein coding | LSC |
| 69 | *rps15* | Protein coding | SSC |
| 70 | *rps16* | Protein coding | LSC |
| 71 | *rps18* | Protein coding | LSC |
| 72 | *rps19* | Protein coding | LSC |
| 73 | *rps2* | Protein coding | LSC |
| 74 | *rps3* | Protein coding | LSC |
| 75 | *rps4* | Protein coding | LSC |
| 76 | *rps7* | Protein coding | IR |
| 77 | *rps8* | Protein coding | LSC |
| 78 | *ycf1* | Protein coding | SSC and IR |
| 79 | *ycf2* | Protein coding | IR |
| 80 | *trnA-UGC* | tRNA | IR |
| 81 | *trnC-GCA* | tRNA | LSC |
| 82 | *trnD-GUC* | tRNA | LSC |
| 83 | *trnE-UUC* | tRNA | LSC |
| 84 | *trnF-GAA* | tRNA | LSC |
| 85 | *trnfM-CAU* | tRNA | LSC |
| 86 | *trnG-GCC* | tRNA | LSC |
| 87 | *trnG-UCC* | tRNA | LSC |
| 88 | *trnH-GUG* | tRNA | LSC |
| 89 | *trnI-CAU* | tRNA | IR |
| 90 | *trnI-GAU* | tRNA | IR |
| 91 | *trnK-UUU* | tRNA | LSC |
| 92 | *trnL-CAA* | tRNA | IR |
| 93 | *trnL-UAA* | tRNA | LSC |
| 94 | *trnL-UAG* | tRNA | SSC |
| 95 | *trnM-CAU* | tRNA | LSC |
| 96 | *trnN-GUU* | tRNA | IR |
| 97 | *trnP-UGG* | tRNA | LSC |
| 98 | *trnQ-UUG* | tRNA | LSC |
| 99 | *trnR-ACG* | tRNA | IR |
| 100 | *trnR-UCU* | tRNA | LSC |
| 101 | *trnS-GCU* | tRNA | LSC |
| 102 | *trnS-GGA* | tRNA | LSC |
| 103 | *trnS-UGA* | tRNA | LSC |
| 104 | *trnT-GGU* | tRNA | LSC |
| 105 | *trnT-UGU* | tRNA | LSC |
| 106 | *trnV-GAC* | tRNA | IR |
| 107 | *trnV-UAC* | tRNA | LSC |
| 108 | *trnW-CCA* | tRNA | LSC |
| 109 | *trnY-GUA* | tRNA | LSC |
| 110 | *rrn16* | rRNA | IR |
| 111 | *rrn23* | rRNA | IR |
| 112 | *rrn4.5* | rRNA | IR |
| 113 | *rrn5* | rRNA | IR |
