## Supplementary material for "A haplotype-resolved, chromosome-scale genome assembly and annotation for *Carya glabra* (pignut hickory; Juglandaceae)": Table S4

Table S4. Annotated genes in the *Carya glabra* mitochondrial genome. The mitogenome includes two chromosomes: mtChr1 and mtChr2.

| No. | Gene | Type | Location | Origin |
|---|---|---|---|---|
| 1 | *atp1* | Protein coding | mtChr1 | Native |
| 2 | *atp4* | Protein coding | mtChr1 | Native |
| 3 | *atp6* | Protein coding | mtChr1 | Native |
| 4 | *atp8* | Protein coding | mtChr1 | Native |
| 5 | *atp9* | Protein coding | mtChr1, mtChr2 | Native |
| 6 | *ccmB* | Protein coding | mtChr1, mtChr2 | Native |
| 7 | *ccmC* | Protein coding | mtChr1 | Native |
| 8 | *ccmFC* | Protein coding | mtChr1 | Native |
| 9 | *ccmFN* | Protein coding | mtChr1, mtChr2 | Native |
| 10 | *cob* | Protein coding | mtChr1 | Native |
| 11 | *cox1* | Protein coding | mtChr1 | Native |
| 12 | *cox2* | Protein coding | mtChr1, mtChr2 | Native |
| 13 | *cox3* | Protein coding | mtChr1 | Native |
| 14 | *matR* | Protein coding | mtChr1, mtChr2 | Native |
| 15 | *mttB* | Protein coding | mtChr1 | Native |
| 16 | *nad1* | Protein coding | mtChr1, mtChr2 | Native |
| 17 | *nad2* | Protein coding | mtChr1, mtChr2 | Native |
| 18 | *nad3* | Protein coding | mtChr1, mtChr2 | Native |
| 19 | *nad4* | Protein coding | mtChr1 | Native |
| 20 | *nad4L* | Protein coding | mtChr1 | Native |
| 21 | *nad5* | Protein coding | mtChr1, mtChr2 | Native |
| 22 | *nad6* | Protein coding | mtChr1 | Native |
| 23 | *nad7* | Protein coding | mtChr1 | Native |
| 24 | *nad9* | Protein coding | mtChr1 | Native |
| 25 | *rpl10* | Protein coding | mtChr1, mtChr2 | Native |
| 26 | *rpl16* | Protein coding | mtChr1 | Native |
| 27 | *rpl2* | Protein coding | mtChr1, mtChr2 | Native |
| 28 | *rpl5* | Protein coding | mtChr1 | Native |
| 29 | *rps1* | Protein coding | mtChr1 | Native |
| 30 | *rps10* | Protein coding | mtChr1, mtChr2 | Native |
| 31 | *rps12* | Protein coding | mtChr1, mtChr2 | Native |
| 32 | *rps14* | Protein coding | mtChr1 | Native |
| 33 | *rps19* | Protein coding | mtChr1 | Native |
| 34 | *rps3* | Protein coding | mtChr1 | Native |
| 35 | *rps4* | Protein coding | mtChr1 | Native |
| 36 | *sdh3* | Protein coding | mtChr1, mtChr2 | Native |
| 37 | *sdh4* | Protein coding | mtChr1 | Native |
| 38 | *petN* | Protein coding | mtChr1 | Plastome |
| 39 | *psaJ* | Protein coding | mtChr1 | Plastome |
| 40 | *psbJ* | Protein coding | mtChr1 | Plastome |
| 41 | *rpl20* | Protein coding | mtChr1 | Plastome |
| 42 | *rpl33* | Protein coding | mtChr1 | Plastome |
| 43 | *rrn18S* | rRNA | mtChr1 | Native |
| 44 | *rrn26S* | rRNA | mtChr1 | Native |
| 45 | *rrn5S* | rRNA | mtChr1 | Native |
| 46 | *trnC-GCA* | tRNA | mtChr1 | Bacteria |
| 47 | *trnC-GCA* | tRNA | mtChr1 | Native |
| 48 | *trnE-UUC* | tRNA | mtChr1 | Native |
| 49 | *trnF-GAA* | tRNA | mtChr1 | Native |
| 50 | *trnfM-CAU* | tRNA | mtChr1 | Native |
| 51 | *trnG-GCC* | tRNA | mtChr1 | Native |
| 52 | *trnI-CAU* | tRNA | mtChr1 | Native |
| 53 | *trnK-UUU* | tRNA | mtChr1, mtChr2 | Native |
| 54 | *trnP-UGG* | tRNA | mtChr1 | Native |
| 55 | *trnQ-UUG* | tRNA | mtChr1 | Native |
| 56 | *trnS-GCU* | tRNA | mtChr1 | Native |
| 57 | *trnS-UGA* | tRNA | mtChr1 | Native |
| 58 | *trnY-GUA* | tRNA | mtChr1 | Native |
| 59 | *trnA-UGC* | tRNA | mtChr1 | Plastome |
| 60 | *trnD-GUC* | tRNA | mtChr1 | Plastome |
| 61 | *trnH-GUG* | tRNA | mtChr1 | Plastome |
| 62 | *trnI-CAU* | tRNA | mtChr1 | Plastome |
| 63 | *trnL-CAA* | tRNA | mtChr1 | Plastome |
| 64 | *trnM-CAU* | tRNA | mtChr1 | Plastome |
| 65 | *trnN-GUU* | tRNA | mtChr1 | Plastome |
| 66 | *trnP-UGG* | tRNA | mtChr1 | Plastome |
| 67 | *trnV-GAC* | tRNA | mtChr1 | Plastome |
| 68 | *trnW-CCA* | tRNA | mtChr1, mtChr2 | Plastome |
