## Supplementary material for "A haplotype-resolved, chromosome-scale genome assembly and annotation for *Carya glabra* (pignut hickory; Juglandaceae)": Table S5

Table S5. Chloroplast-derived segments in the *Carya glabra* mitochondrial genome. *Fragmentary gene; +gene containing premature stop codon.

| Location | Aligned length (bp) | Identity to plastome (%) | Plastome gene(s) |
|---|---|---|---|
| mtChr1: 335851-350810 | 15,031 | 99.2 | *trnA-UGC*, *trnI-CAU*, *trnL-CAA*, *trnV-GAC*, *rps12**, *rpl23**, *rrn16S**, *ndhB*+, *rps7*+, *ycf2*+ |
| mtChr1: 366061-367918 | 2,137 | 84.3 | *psaJ*, *rpl20*, *rpl33*, *rps18*+ |
| mtChr1: 288398-289764 | 1,379 | 99.0 | *rpl2**, *rpl23** |
| mtChr1: 71318-72550 | 1,233 | 71.4 | *trnP-UGG*, *trnW-CCA*, *petG*+, *petL*+ |
| mtChr1: 441717-442798 | 1,083 | 94.7 | *trnD-GUC* |
| mtChr1: 367903-368823 | 936 | 97.1 | *petN* |
| mtChr1: 102562-103322 | 769 | 92.9 | *psaA**, *psaB** |
| mtChr1: 72510-73190 | 681 | 99.7 | *rpoC2** |
| mtChr1: 249552-249976 | 425 | 99.5 | *trnI-GAU** |
| mtChr1: 357130-357276 | 147 | 90.5 | *psbJ* |
| mtChr1: 283627-283714 | 88 | 95.5 | *trnH-GUG* |
| mtChr1: 474776-474858 | 83 | 100.0 | *trnN-GUU* |
| mtChr1: 153840-153917 | 78 | 94.9 | *trnM-CAU* |
| mtChr1: 376921-376997 | 77 | 84.4 | *trnM-CAU* |
