## Supplementary material for "A haplotype-resolved, chromosome-scale genome assembly and annotation for *Carya glabra* (pignut hickory; Juglandaceae)": Table S6

Table S6. Omni-C library quality control report from Phase Genomics' hic_qc pipeline.

| Category | Omni-C library statistics | Expected values |
|---|---|---|
| Percentage of read pairs spanning at least 10 kb among all high-quality read pairs mapped to the same unitig of at least 10 kb in length | 6.7% | >3.0% |
| Percentage of high-quality read pairs mapping to two different unitigs, each greater than 10 kb in length | 7.1% | >2.5% |

Note: high-quality read pairs are defined as those with a minimum mapping quality of 20, maximum edit distance of 5, and no duplicate reads.
