## Supplementary material for "A haplotype-resolved, chromosome-scale genome assembly and annotation for *Carya glabra* (pignut hickory; Juglandaceae)": Table S7

Table S7. Lengths (in Mb) of the 64 assembled pseudo-chromosomes of *Carya glabra*.

| Pseudo-chromosome | Haplotype A | Haplotype B | Haplotype C | Haplotype D |
|---|---|---|---|---|
| 1 | 53.0 | 52.6 | 52.1 | 49.6 |
| 2 | 36.2 | 35.2 | 35.2 | 34.3 |
| 3 | 50.2 | 48.6 | 47.3 | 47.2 |
| 4 | 37.5 | 36.7 | 36.2 | 36.2 |
| 5 | 45.4 | 43.7 | 43.0 | 42.0 |
| 6 | 31.5 | 30.6 | 30.6 | 30.3 |
| 7 | 39.6 | 39.4 | 37.8 | 36.9 |
| 8 | 33.6 | 32.9 | 32.0 | 31.1 |
| 9 | 42.3 | 40.5 | 39.9 | 39.8 |
| 10 | 33.8 | 33.5 | 33.1 | 32.9 |
| 11 | 41.2 | 41.1 | 39.6 | 38.8 |
| 12 | 24.9 | 24.7 | 23.4 | 20.9 |
| 13 | 32.0 | 31.9 | 31.6 | 28.8 |
| 14 | 30.5 | 29.4 | 29.2 | 28.5 |
| 15 | 41.4 | 37.7 | 36.8 | 35.4 |
| 16 | 27.5 | 26.9 | 26.8 | 26.7 |
