## Supplementary material for "A haplotype-resolved, chromosome-scale genome assembly and annotation for *Carya glabra* (pignut hickory; Juglandaceae)": Table S8

Table S8. Summary of repetitive element annotation in *Carya glabra*.

| Category | | Hap. A | Hap. B | Hap. C | Hap. D |
|---|---|---|---|---|---|
| Transposable elements | Class I: retrotransposons | 27.2% | 27.1% | 24.7% | 26.9% |
| | Class II: DNA transposons | 19.5% | 19.9% | 21.5% | 19.4% |
| Simple repeats | | 1.2% | 1.2% | 1.3% | 1.3% |
| Low complexity | | 0.2% | 0.2% | 0.2% | 0.2% |
| Unclassified interspersed repeats | | 6.9% | 6.1% | 6.2% | 6.0% |
| Total | | 55.0% | 54.4% | 54.0% | 53.8% |
