## Supplementary material for "A haplotype-resolved, chromosome-scale genome assembly and annotation for *Carya glabra* (pignut hickory; Juglandaceae)": Table S9

Table S9. Statistics of finalized gene models predicted for four haplotypes from *Carya glabra*.

| Haplotype | A | B | C | D |
|---|---|---|---|---|
| Gene no. | 30,947 | 31,087 | 30,369 | 30,110 |
| Average gene length (bp) | 4,398 | 4,364 | 4,460 | 4,381 |
| Average CDS length (bp) | 1,241 | 1,239 | 1,254 | 1,240 |
| Average exons per gene | 5.0 | 5.0 | 5.0 | 5.0 |
| BUSCO (%) | 97.7 | 97.1 | 96.5 | 94.9 |
| Genes assigned to gene families (%) | 94.3 | 93.9 | 94.7 | 94.4 |
| Genes annotated with GO terms (%) | 79.7 | 79.4 | 79.9 | 79.4 |
| Genes annotated with protein domains (%) | 86.3 | 85.9 | 86.5 | 86.3 |
| Core gene family completeness | 0.982 | 0.982 | 0.978 | 0.967 |

Note: the statistics are based on the longest isoform of each gene. GO: gene ontology.
