## Supplementary material for "A haplotype-resolved, chromosome-scale genome assembly and annotation for *Carya glabra* (pignut hickory; Juglandaceae)": Table S10

Table S10. Four major classes of plant disease resistance genes (*R* genes) identified in *Carya glabra* and three other *Carya* species with assembled genomes.

| *R* gene class | *C. glabra* | | | | *C. illinoinensis*[1] | *C. sinensis*[2] | *C. cathayensis*[2] |
|---|---|---|---|---|---|---|---|
| | Hap. A | Hap. B | Hap. C | Hap. D | | | |
| CNL | 56 | 61 | 59 | 41 | 70 | 44 | 29 |
| TNL | 39 | 83 | 96 | 34 | 65 | 11 | 26 |
| RLP | 214 | 192 | 189 | 226 | 217 | 300 | 375 |
| RLK | 316 | 302 | 311 | 307 | 372 | 330 | 370 |
| Total | 625 | 638 | 655 | 608 | 724 | 685 | 800 |

Note: the CNL protein contains the coiled-coil domain, the nucleotide-binding site (NBS) domain, and the leucine-rich repeat (LRR) domain. The TNL protein contains the Toll-interleukin receptor-like domain, the NBS domain, and the LRR domain. The RLP (receptor-like protein) protein contains the transmembrane (TM) domain and the LRR domain. The RLK (receptor-like kinase) protein contains the TM domain, the LRR domain, and the kinase domain. [1]The statistics are from *C. illinoinensis* cv. 'Pawnee' (Lovell et al. 2021). [2]Both genomes are from Zhang et al. 2024b. Hap.: haplotype.
