## Supplementary material for "A haplotype-resolved, chromosome-scale genome assembly and annotation for *Carya glabra* (pignut hickory; Juglandaceae)": Table S11

Table S11. Putative *Carya glabra* plant disease resistance genes (*R* genes) identified in the syntenic regions corresponding to the major quantitative trait locus (QTL) associated with phylloxera resistance in *Carya illinoinensis*.

| Haplotype | Gene name | *R* gene class | Location |
|---|---|---|---|
| A | dhCarGlab.v1a1.Chr16A.g01360.t1 | TNL | Chr. 16A: 1973676 - 1977343 |
| A | dhCarGlab.v1a1.Chr16A.g01410.t1 | TNL | Chr. 16A: 2185240 - 2190125 |
| A | dhCarGlab.v1a1.Chr16A.g01420.t1 | RLP | Chr. 16A: 2268899 - 2269806 |
| A | dhCarGlab.v1a1.Chr16A.g01430.t1 | TNL | Chr. 16A: 2270798 - 2275685 |
| A | dhCarGlab.v1a1.Chr16A.g01450.t1 | TNL | Chr. 16A: 2291459 - 2295800 |
| A | dhCarGlab.v1a1.Chr16A.g01460.t1 | TNL | Chr. 16A: 2325172 - 2331391 |
| A | dhCarGlab.v1a1.Chr16A.g01480.t1 | TNL | Chr. 16A: 2412331 - 2417424 |
| A | dhCarGlab.v1a1.Chr16A.g01500.t1 | RLK | Chr. 16A: 2433195 - 2436628 |
| B | dhCarGlab.v1a1.Chr16B.g01550.t1 | TNL | Chr. 16B: 2162361 - 2165884 |
| B | dhCarGlab.v1a1.Chr16B.g01560.t2 | TNL | Chr. 16B: 2199479 - 2204805 |
| B | dhCarGlab.v1a1.Chr16B.g01570.t1 | TNL | Chr. 16B: 2271505 - 2275301 |
| B | dhCarGlab.v1a1.Chr16B.g01600.t2 | TNL | Chr. 16B: 2291370 - 2296652 |
| B | dhCarGlab.v1a1.Chr16B.g01630.t2 | TNL | Chr. 16B: 2357117 - 2363523 |
| B | dhCarGlab.v1a1.Chr16B.g01640.t2 | RLP | Chr. 16B: 2370716 - 2371754 |
| B | dhCarGlab.v1a1.Chr16B.g01650.t1 | TNL | Chr. 16B: 2371918 - 2375633 |
| B | dhCarGlab.v1a1.Chr16B.g01680.t1 | TNL | Chr. 16B: 2444556 - 2449470 |
| B | dhCarGlab.v1a1.Chr16B.g01700.t1 | TNL | Chr. 16B: 2468763 - 2473767 |
| B | dhCarGlab.v1a1.Chr16B.g01770.t1 | RLK | Chr. 16B: 2535654 - 2539083 |
| C | dhCarGlab.v1a1.Chr16C.g01400.t1 | TNL | Chr. 16C: 2136794 - 2140456 |
| C | dhCarGlab.v1a1.Chr16C.g01410.t2 | TNL | Chr. 16C: 2153782 - 2158982 |
| C | dhCarGlab.v1a1.Chr16C.g01440.t1 | TNL | Chr. 16C: 2386033 - 2391055 |
| C | dhCarGlab.v1a1.Chr16C.g01480.t1 | TNL | Chr. 16C: 2497010 - 2502078 |
| C | dhCarGlab.v1a1.Chr16C.g01510.t1 | TNL | Chr. 16C: 2547531 - 2553573 |
| C | dhCarGlab.v1a1.Chr16C.g01530.t1 | TNL | Chr. 16C: 2660559 - 2665364 |
| C | dhCarGlab.v1a1.Chr16C.g01550.t1 | TNL | Chr. 16C: 2738972 - 2742700 |
| C | dhCarGlab.v1a1.Chr16C.g01560.t1 | TNL | Chr. 16C: 2743843 - 2748369 |
| C | dhCarGlab.v1a1.Chr16C.g01600.t1 | TNL | Chr. 16C: 2835169 - 2841464 |
| C | dhCarGlab.v1a1.Chr16C.g01680.t1 | TNL | Chr. 16C: 2982302 - 2988728 |
| C | dhCarGlab.v1a1.Chr16C.g01700.t1 | RLK | Chr. 16C: 3025435 - 3028868 |
| D | dhCarGlab.v1a1.Chr16D.g01310.t1 | TNL | Chr. 16D: 1961595 - 1966668 |
| D | dhCarGlab.v1a1.Chr16D.g01320.t1 | TNL | Chr. 16D: 1986524 - 1991236 |
| D | dhCarGlab.v1a1.Chr16D.g01390.t1 | TNL | Chr. 16D: 2323753 - 2330174 |
| D | dhCarGlab.v1a1.Chr16D.g01420.t1 | RLP | Chr. 16D: 2521824 - 2522679 |
| D | dhCarGlab.v1a1.Chr16D.g01430.t1 | TNL | Chr. 16D: 2523457 - 2528122 |
| D | dhCarGlab.v1a1.Chr16D.g01440.t1 | TNL | Chr. 16D: 2545456 - 2549333 |
| D | dhCarGlab.v1a1.Chr16D.g01490.t1 | TNL | Chr. 16D: 2650932 - 2655992 |
| D | dhCarGlab.v1a1.Chr16D.g01520.t1 | RLK | Chr. 16D: 2694631 - 2698064 |

Note: the TNL protein contains the Toll-interleukin receptor-like domain, the nucleotide-binding site (NBS) domain, and the leucine-rich repeat (LRR) domain. The RLP (receptor-like protein) protein contains the transmembrane (TM) domain and the LRR domain. The RLK (receptor-like kinase) protein contains the TM domain, the LRR domain, and the kinase domain. Chr.: chromosome.
