## Supplementary material for "A haplotype-resolved, chromosome-scale genome assembly and annotation for *Carya glabra* (pignut hickory; Juglandaceae)": Table S12

Table S12. Misannotated and missing genes in previously published *Carya glabra* chloroplast genomes.

| Gene | Published *Carya glabra* chloroplast genomes on GenBank | | |
|---|---|---|---|
| | BK061156 (Luo et al. 2021) | OR099205 (Liu et al. 2025) | NC_067504 (Xi et al. 2022) |
| *atpB* | | Missing | |
| *infA* | | | Missing |
| *petB* | | | Exon 1 missing |
| *petD* | | | Exon 1 missing |
| *psbB* | | Misannotated as *psi* | |
| *psbZ* | | Misannotated as *lhbA* | |
| *rpl16* | | | Exon 1 missing |
| *rpoB* | | Missing | |
| *ycf15* | Included | Included | Included |
| *trnA-UGC* | Two additional copies misannotated at 105,498–105,557 and 144,984–145,043 | | Two additional copies misannotated at 105,495–105,554 and 144,981–145,040 |
| *trnK-UUU* | One additional copy misannotated at 6,403–6,462 | | Misannotated at 6,400–6,459 |
| *tnrM-CAU* | Two additional copies misannotated at 91,947–92,021 and 158,520–158,594 | One additional copy misannotated at 35,273–35,331 | |
| *trnP-GGG* | | Misannotated at 72,042–72,112 | |
| *trnT-GGU* | | One additional copy misannotated at 57,114–57,172 | |
| *rrn16* | | Missing | |
| *rrn23* | | Missing | |
| *rrn4.5* | | Missing | |
| *rrn5* | | Missing | |
